# Long-range neuropeptide relay as a central-peripheral communication mechanism for the context-dependent modulation of interval timing behaviors

**DOI:** 10.1101/2024.06.03.597273

**Authors:** Tianmu Zhang, Zekun Wu, Yutong Song, Wenjing Li, Yanying Sun, Xiaoli Zhang, Kyle Wong, Justine Schweizer, Khoi-Nguyen Ha Nguyen, Alex Kwan, Woo Jae Kim

## Abstract

Neuropeptides play crucial roles in regulating context-dependent behaviors, but the underlying mechanisms remain elusive. We investigate the role of the neuropeptide SIFa and its receptor SIFaR in regulating two distinct mating duration behaviors in male *Drosophila*: Longer-Mating-Duration (LMD) and Shorter-Mating-Duration (SMD). We found that SIFaR expression in specific neurons is required for both LMD and SMD behaviors. Social context and sexual experience lead to synaptic reorganization between SIFa and SIFaR neurons, altering internal states of brain. We revealed that the SIFa-SIFaR/Crz-CrzR neuropeptide relay pathway is essential for generating distinct interval timing behaviors, with Crz neurons being responsive to the activity of SIFa neurons. Additionally, CrzR expression in non-neuronal cells is critical for regulating LMD and SMD behaviors. Our study provides insights into how neuropeptides and their receptors modulate context-dependent behaviors through synaptic plasticity and calcium signaling, with implications for understanding the neural circuitry underlying interval timing and neuropeptidergic system modulation of behavioral adaptations.

## INTRODUCTION

Plasticity in animal behavior is dependent on the capacity to integrate external stimuli and internal states from a fluctuating environment and, as a result, to modify activity in neuronal circuits of the brain^1,2^. In *Drosophila*, many of the neural systems that are known to influence behavioral circuits in a state-dependent manner and create switches between behaviors make use of various neuropeptides or biogenic amines like serotonin, dopamine, or octopamine^3–9^. These neuromodulatory circuits or pathways are not always hardwired; they depend on paracrine signaling or volume transmission, which is based on non-synaptic release of an amine or neuropeptide^4,10^. Interorgan communication is also represented by neuromodulatory signaling, which is even hormonal and conducted via the circulation^11–13^.

The modulating energy homeostasis between the gut and the brain is one of the most intensively studied neuro-modulatory circuits via the neuronal relay of neuropeptides^13–18^. The hypothalamus in the mammalian brain regulates hunger and satiety by neuropeptide and hormonal relay signaling between the stomach, adipose tissue, and pancreas^13,19^. The relayed event of non-synaptic and diffuse neurotransmission through neuropeptides and their receptors is also essential for the maintenance of metabolic homeostasis in *Drosophila*^1,14,20–24^. However, neuropeptidergic relays within the central nervous system and their detailed circuits modulating complex decision-making behavior are poorly understood.

The neuropeptide SIFa is a neuromodulator that demonstrates plasticity on a molecular and behavioral level. SIFa is expressed by four neurons located in the pars intercerebrails (PI) and they are the most widely arborizing peptidergic neurons in *Drosophila*^25^. SIFa has significant impact on various behaviors including^26,27^, courtship^28,29^, sleep^30–32^, memory^33^, and interval timing^34,35^. SIFa neurons integrate information through several inputs and form a hub to orchestrate many behaviors. The SIFa neurons receive inputs from peptidergic neurons expressing hugin-pyrokinin (PK)/myoinhibitory peptide (MIP) that mediate hunger/satiety^26^, *Drosophila* insulin-like peptides (DILPs) that mediates feeding: fasting rhythm^27^ and sleep^30,31^. Thus, it is anticipated that the SIFa neurons would play a crucial role in the context-dependent orchestration of multiple behaviors^4^.

However, the mechanism by which SIFa signaling controls various behaviors through its receptor SIFaR is not well known, because it is difficult to resolve using ordinary molecular genetics approaches and typical neural connectomics may not necessarily predict peptidergic circuits. Here, we report that SIFa controls two alternate interval timing behaviors through long-range neuropeptide relay signaling by SIFaR and other important neuropeptides and transmits the internal states of the male brain into decision making.

## RESULTS

### Two distinct interval timing behaviors are governed by the adult-specific expression of SIFaR

We have described two distinct male *Drosophila melanogaster* behaviors as a model for investigating the neural circuit principles that determine interval timing. Male fruit flies exhibit the Longer-Mating-Duration (LMD) behavior, which is characterized by a prolonged duration of mating after exposure to competing males. This behavior is believed to be an adaptation to male mating competition (Supplementary information, Fig. S1A)^36–44^. Shorter-Mating-Duration (SMD) is a behavior in which sexually experienced males manifest a shorter duration of mating. This behavior is believed to be an adaptation that allows male flies to conserve energy by mating quickly then continuing to other activities (Supplementary information, Fig. S1B)^45–48^.

We discovered through neuropeptide RNAi screening that neuropeptide SIFa modulates two distinct interval timing behaviors by recording and orchestrating the male’s internal states^49^. The SIFaR, the receptor for SIFa, has been linked to circadian rhythm, feeding, courtship, sleep, and memory extinction; however, its functional properties remain poorly understood^26–28,30,33,50,51^. When we used a pan-neuronal *elav^c155^* driver to knock down *SIFaR* in a neuronal population, both LMD and SMD were disrupted (Fig. 1A, B). To ensure that RNAi did not have an off-target effect, we tested three independent RNAi strains and found that all three RNAi successfully disrupted LMD/SMD when expressed in neuronal populations. (Supplementary information, Fig. S1C-H). We chose to use the HMS00299 line as *SIFaR-RNAi* for all our experiments because it efficiently disrupts LMD/SMD without *UAS-dicer* expression. When we used *tub-GAL80^ts^* to knock down SIFaR in all neuronal populations in an adult-specific manner, both LMD and SMD were also impaired compared to temperature control (Compare Fig. 1E, F; Supplementary information, Fig. S1I, J). SIFaR knockdown in glial cells, on the other hand, has no impact on LMD/SMD behavior (Fig. 1C, D), indicating that SIFaR expression in adult neurons not glia is specifically involved in interval timing behaviors.

**Figure 1.**
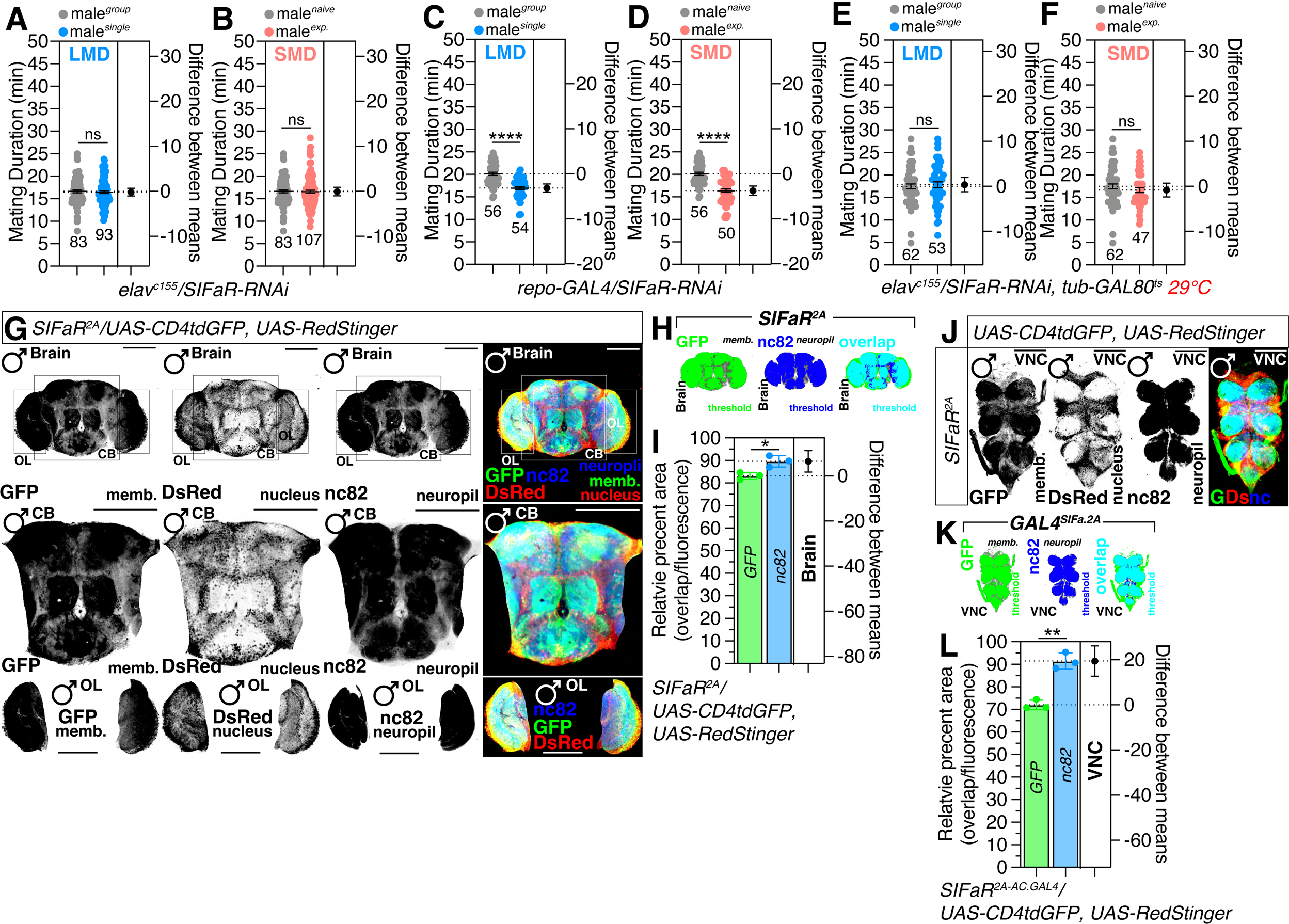
Interval timing is regulated by adult-specific SIFaR-positive neuronal cells. (A-D) LMD and SMD assays for *elav^c155^* and *repo-GAL4* mediated knockdown of SIFaR *via SIFaR-RNAi.* Light grey dots represent naïve males, blue dots represent single reared males and pink dots represent experienced males. Dot plots represent the MD of each male fly. The mean value and standard error are labeled within the dot plot (black lines). Asterisks represent significant differences, as revealed by the Student’s *t* test and ns represents non-significant difference (*\*p<0.05, **p<0.01, ***p< 0.001, ****p< 0.0001*). For detailed methods, see the MATERIALS AND METHODS for a detailed description of the mating duration assay used in this study. (E-F) LMD and SMD assays for *elav^c155^*–mediated knockdown of SIFaR *via SIFaR-RNAi* together with *tub-GAL80^ts^* in 29. (G) Brain of male flies expressing *SIFaR^2A^* together with *UAS-CD4tdGFP, UAS-Redstinger* were immunostained with anti-GFP (green), anti-DsRed (red), and nc82 (blue) antibodies. Scale bars represent 100 μm. Boxes indicate the magnified regions of interest presented in the middle and bottom panels. The left three panels are presented as a grey scale to clearly show the expression patterns of neurons in brain labeled by *SIFaR^2A^* driver. For detailed methods, see the MATERIALS AND METHODS for a detailed description of the immunostaining procedure used in this study. (H) The GFP fluorescence (green, left panel), nc82 (blue, middle panel) and overlapping area of GFP and nc82 (cyan, right panel) in male fly brain was processed using ImageJ software, where a threshold function was applied to distinguish fluorescence from the background. (I) Colocalization analysis of GFP and nc82 staining, normalized to total GFP and nc82 areas. Bars represent the mean GFP (green column) and nc82 (blue column) fluorescence level with error bars representing SEM. Asterisks represent significant differences, as revealed by the Student’s *t* test and ns represents non-significant difference (*\*p<0.05, **p<0.01, ***p< 0.001, ****p< 0.0001*). The same symbols for statistical significance are used in all other figures. See the MATERIALS AND METHODS for a detailed description of the colocalization analysis used in this study. (J) VNC of male flies expressing *SIFaR^2A^* together with *UAS-CD4tdGFP, UAS-Redstinger*. Scale bars represent 100 μm. (K) The GFP fluorescence (green, left panel), nc82 (blue, middle panel) and overlapping area of GFP and nc82 (cyan, right panel) in male fly VNC was processed using ImageJ software, where a threshold function was applied to distinguish fluorescence from the background. (L) Colocalization analysis of GFP and nc82 staining, normalized to total GFP and nc82 areas. See the MATERIALS AND METHODS for a detailed description of the colocalization analysis used in this study. See also Figure S1.

We used the newly developed chemoconnectome (CCT) knock-in *SIFaR-GAL4* (*SIFaR^2A-AC.GAL4^, SIFaR^2A^* hereafter) strain^52^ to examine SIFaR expression in the nervous system, and our results showed that SIFaR is expressed in abundance in the most brain and VNC area (Fig. 1G-J). About 1150 SIFaR^+^ cells are found in the brain, while only 300 are found in the VNC (Supplementary information, Fig. S1K, L). About 620 of the 1150 brain cells are found in the central brain (CB), while 530 are found in the optic lobe (OL) (Supplementary information, Fig. S1M, N). Area of nc82-labeled neuropil covered by SIFaR^+^ cell membrane is nearly identical in brain, VNC, CB, and OL (Supplementary information, Fig. S1O-R). Neuropil is a synaptic dense region in the nervous system composed of mostly unmyelinated axons, dendrites and glial cell processes^53^. Monoclonal antibody nc82 identifies *Drosophila* Bruchpilot (Brp), a CAST1/ERC family member and active zone component. Thus, if the membrane marker of certain neuronal populations strongly overlaps with the neuropil marker nc82, it could imply that the membrane of those neuronal populations is also present in the active zone^54^. We found that about 83% of brain and 71% of VNC SIFaR^+^ neurons overlap with nc82-positive active zones of neuropil (Fig. 1H, I; Fig. 1K, L). All these findings indicate that neuronal expression of SIFaR is essential for inducing interval timing behaviors.

### SIFaR expression in specific cell populations is both required and sufficient to maintain interval timing behaviors

To identify the minimal region labeling GAL4 drivers for testing the functionality of SIFaR-mediated interval timing behaviors, we examined four custom GAL4 drivers^55^ targeting the SIFaR promoter region (Fig. 2A). LMD and SMD disappeared when SIFaR-RNAi was co-expressed with *GAL4^57F10^* and *GAL4^24F06^*, indicating that SIFaR expression in these GAL4 driver-expressing cells is required for interval timing behaviors (Fig. 2B-I).

**Figure 2.**
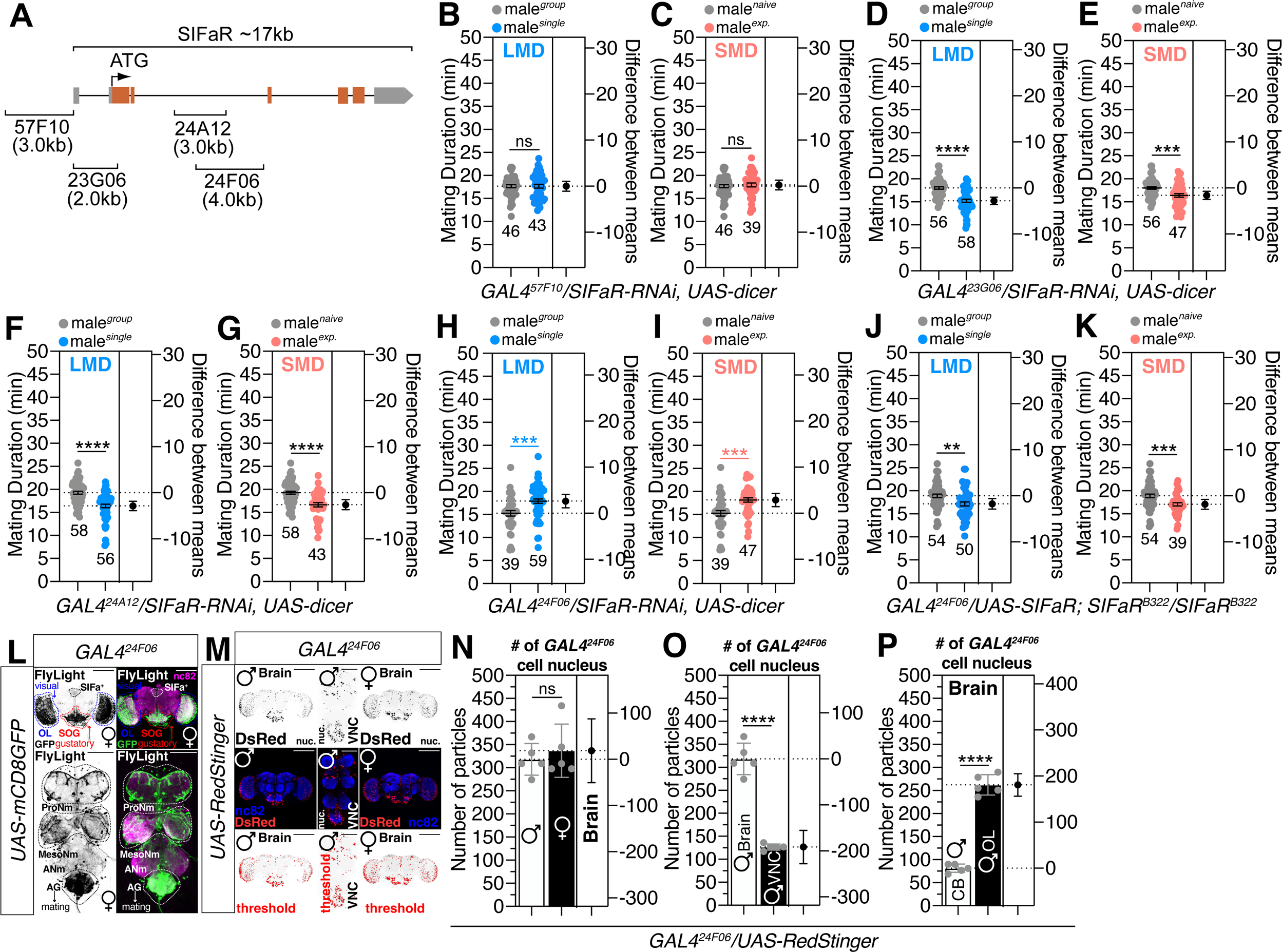
The expression of *SIFaR^24F06^* neurons is necessary and sufficient for LMD/SMD. (A) Diagram of the captured promoter region within each *SIFaR-GAL4* construct. (B-I) LMD and SMD assays of (B-C) *GAL4^57F10^*, (D-E) *GAL4^23G06^*, (F-G) *GAL4^24A12^* and (H-I) *GAL4^24F06^* mediated knockdown of SIFaR *via SIFaR-RNAi* together with *UAS-dicer*. (J-K) Genetic rescue experiments of LMD and SMD assays for *GAL4* mediated overexpression of SIFaR *via GAL4^24F06^* in *SIFaR* mutant background flies. (L) Brain and VNC of male flies expressing *GAL4^24F06^* together with *UAS-mCD8GFP* in Flylight. Scale bars represent 100 μm in brain panels and 50 μm in VNC panels. Dashed circles indicate the region of interest. (M) Brain and VNC of male and female flies expressing *GAL4^24F06^* together with *UAS-RedStinger*. Scale bars represent 100 μm in brain panels and 50 μm in VNC panels. (N-P) Quantification of cell number. The conditions of flies are described above: male, female fly brain; male fly brain and VNC; male fly CB and OL. Bars represent the mean particle number with error bars representing SEM. Asterisks represent significant differences, as revealed by the Student’s *t* test and ns represents non-significant difference (*\*p<0.05, **p<0.01, ***p< 0.001, ****p< 0.0001*). See the MATERIALS AND METHODS for a detailed description of the particle analysis used in this study. See also Figures S2 and S3.

To determine the cell populations in which SIFaR expression is sufficient to maintain interval timing behaviors, we conducted genetic rescue experiments with the *SIFaR^B322^* homozygous lethal mutant strain. As shown in Table 1, the expression of *UAS-SIFaR* with *GAL4^57F10^* and *GAL4^23G06^* drivers in a *SIFaR^B322^* mutant background cannot rescue the lethality of the mutant, whereas *GAL4^24A12^* and *GAL4^24F06^* can rescue the lethality of the *SIFaR^B322^* mutant by expressing *UAS-SIFaR*. Among these two GAL4 drivers, only *GAL4^24F06^* can rescue both lethality and interval timing behaviors in a *SIFaR^B322^* mutant background by expressing *UAS-SIFaR* (Fig. 2J, K), indicating that the cells labeled by *GAL4^24F06^* driver contain neurons that are critical for both lethality and interval timing behaviors via SIFa-SIFaR neuropeptidergic signaling.

**Table 1.** Summary of *SIFaR^B322^* mutant genetic rescue data.

| **GAL4** | **Genotype** | **Lethality**<br>**(non-TM6B/TM6B)** | **Courtship** | **LMD** | **SMD** |
| --- | --- | --- | --- | --- | --- |
| *57F10* | *GAL4*<sup>57F10</sup>-, *SIFaR*<sup>B322</sup><br>/*UAS-SIFaR*; *SIFaR*<sup>B322</sup> | 0/334 | N/A | N/A | N/A |
| *23G06* | *GAL4*<sup>23G06</sup>, *SIFaR*<sup>B322</sup><br>/*UAS-SIFaR*; *SIFaR*<sup>B322</sup> | 0/170 | N/A | N/A | N/A |
| *24A12* | *GAL4*<sup>24A12</sup>, *SIFaR*<sup>B322</sup><br>/*UAS-SIFaR*; *SIFaR*<sup>B322</sup> | 80/90 | no successful<br>mating | N/A | N/A |
| *24F06* | *GAL4*<sup>24F06</sup>, *SIFaR*<sup>B322</sup><br>/*UAS-SIFaR*; *SIFaR*<sup>B322</sup> | 81/90 | normal | normal | normal |

The expression patterns (Fig. 2L) and the number of *GAL4^24F06^* expressing cells (∼320 cells in both sexes, Fig. 2M, N) are similar between male and female brains, indicating that the critical cell population for both lethality and interval timing behaviors via SIFaR signaling exists in both sexes. The number of *GAL4^24F06^* expressing cells is substantially lower in the VNC than in the brain (∼125 cells in male VNC; Fig. 2O). In the brain, the central brain (CB) contains approximately 75 cells and the optic lobe (OL) contains approximately 260 cells (Fig. 2P). All of these results indicate that *GAL4^24F06^* labeled cells are highly expressed in the brain and VNC of both sexes.

### Neurons expressing essential SIFaR form functional synapses with SIFa neurons in social context-dependent manner

To compare the expression patterns of *GAL4^24F06^*-positive neurons with those of all other SIFaR-expressing cells, we utilized genetic intersectional methods with a recently developed SIFaR knock-in line, *SIFaR^2A^* driver (Supplementary information, Fig. S2A)^52^. Approximately 87% and 80% of *GAL4^24F06^*-positive neurons in the brain and VNC overlap with SIFaR^2A^-positive cells, respectively (Supplementary information, Fig. S2B, E). Among *GAL4^24F06^* neurons in the brain, OL neurons have more overlap with *SIFaR^2A^* neurons than CB neurons (Supplementary information, Fig. S2C, D). Among VNC *GAL4^24F06^*-positive neurons, AG neurons overlap significantly with *SIFaR^2A^*-positive neurons relative to the ventral association center (VAC) (Supplementary information, Fig. S2F, G). These findings indicate that the newly identified *GAL4^24F06^* neurons are pertinent SIFaR-positive cells, with particularly high expression in OL, SOG, and AG neurons.

To determine the neuronal architectures between SIFa neurons and the identified essential SIFaR-expressing neurons, we labeled both SIFa and SIFaR neurons using genetic intersectional methods^56^. As previously reported, SIFa neurons arborize extensively throughout the CNS, but the neuronal processes of *GAL4^24F06^*-positive neurons are enriched in the optic lobe (OL), sub-esophageal ganglion (SOG), and abdominal ganglion (AG) (GFP signal in Fig. 2L). Neuronal processes that are positive for SIFa and SIFaR strongly overlap in the prow (PRW), prothoracic and metathoracic neuromere (ProNm and MesoNm), and AG regions (yellow signals in supplementary information, Fig. S3A). We quantified these overlapping neuronal processes between SIFa- and SIFaR-positive neurons and found that approximately 18% of SIFa neurons and 52% of *GAL4^24F06^*-positive neurons overlap in brain (Supplementary information, Fig. S3B, C), whereas approximately 48% of SIFa and 54% of *GAL4^24F06^*-positive neurons overlap in VNC (Supplementary information, Fig. S3D, E). These findings suggest that SIFa neurons and *GAL4^24F06^*-positive neurons form more synapses in the VNC than in the brain.

Next, we characterized the postsynaptic (dendritic) and presynaptic (synaptic terminal) neuronal processes of SIFaR^24F06^-positive neurons by using DenMark^57^ and synaptotagmin-GFP (syt.eGFP)^58^. Dendrites of *GAL4^24F06^*-positive neurons were found to be enriched in SOG, ProNm, and AG regions (dendrites panels and red signals in Supplementary information, Fig. S3F), whereas synaptic terminals of these neurons were found to be widely distributed throughout the CNS (presynaptic terminals panels and green signals in Fig. 3F). The overlap region of DenMark and syt.eGFP signals was highly enriched in both SOG and ProNm regions, indicating that these regions are where *GAL4^24F06^* neurons form interconnected networks (yellow signals in supplementary information, Fig. S3F). We also discovered that approximately 65% and 55% of DenMark signals overlap with syt.eGFP in the brain and VNC, respectively, whereas only 10% and 4% of syt.eGFP signals overlap with DenMark signals (Supplementary information, Fig. S3G-J). These results imply that *GAL4^24F06^*-positive neurons are interconnected in specific regions of the brain and the VNC (Supplementary information, Fig. S3K).

**Figure 3.**
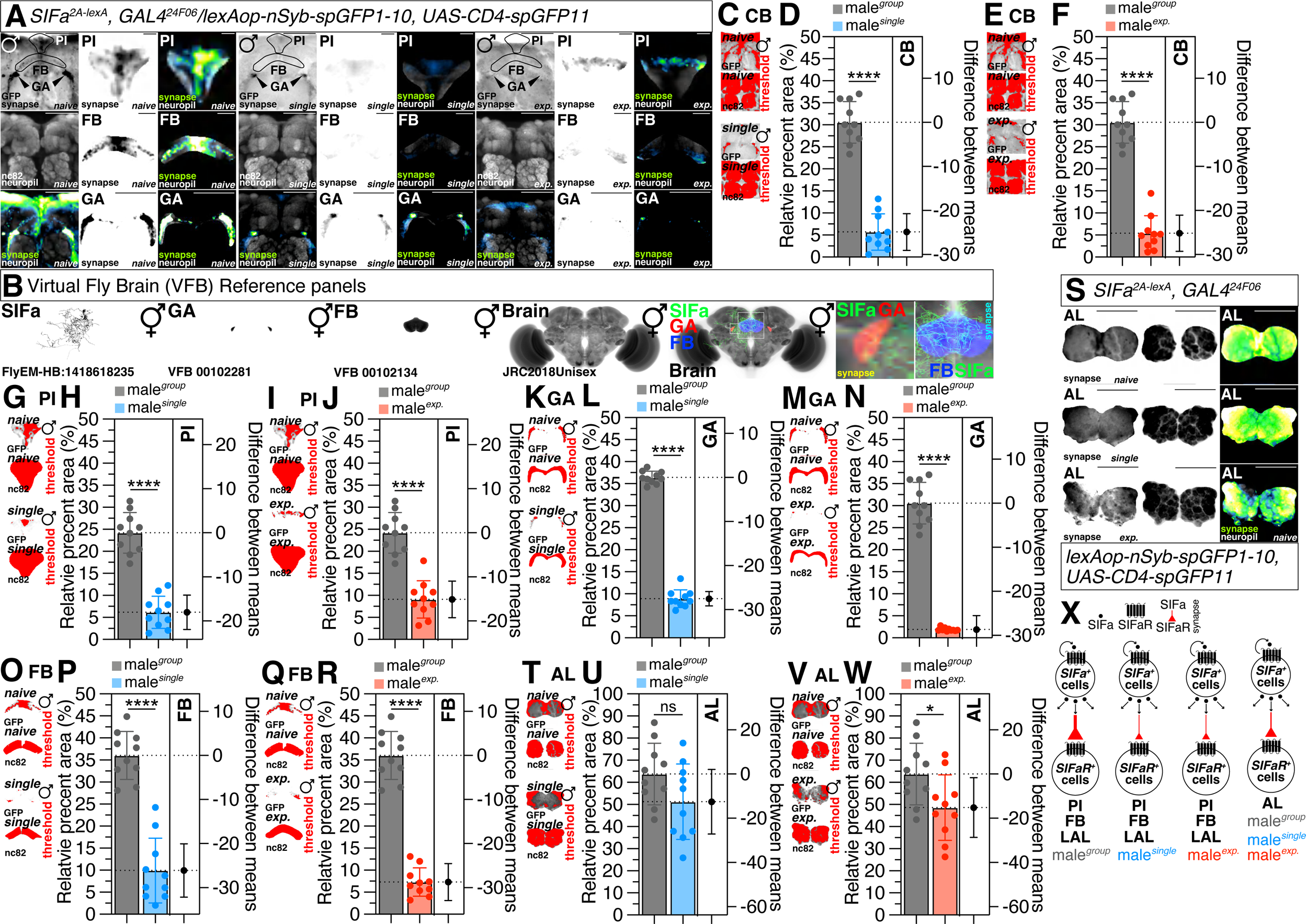
Social context modulates the formation of synapses between *SIFa^2A-lexA^* and *GAL4^24F06^* neurons. (A) GRASP assay for *SIFa^2A-lexA^* and *GAL4^24F06^* in PI region (upper panels), FB (middle panels) and GA region (bottom panels) of naïve (left three columns), single (middle three columns) and experienced (right three columns) male flies. Male flies expressing *SIFa^2A-lexA^, GAL4^24F06^* and *lexAop-nsyb-spGFP1-10, UAS-CD4-spGFP11* were dissected after 5 days of growth (mated male flies had 1-day of sexual experience with virgin females). Brains of male flies were immunostained with anti-GFP (green) and anti-nc82 (blue) antibodies. The middle panels in each condition are presented as a grey scale to clearly show the synapses connection between *SIFa^2A-lexA^* and *GAL4^24F06^*. Circles indicate the regions of interest presented in the panels besides. Synaptic transmission occurs from *SIFa^2A-lexA^* to *GAL4^24F06^*. Scale bars represent 50 μm in left columns in each condition, 10 μm PI panels, 25 μm in FB panels and 50 μm in GA panels. GFP is pseudo-colored as “Green fire blue”. (B) Expression pattern of *SIFa* in Virtual Fly Brain (VFB) and co-localization between *SIFa* and GA/FB. (C-R) Quantification of relative value for synaptic area which are formed between *SIFa^2A-lexA^* and *GAL4^24F06^* in (C-D) CB, (G-H) PI, (K-L) GA and (O-P) FB between naïve and single male flies. The same quantification was performed for the relative synaptic area in these brain regions between naïve and experienced male flies. The synaptic interactions were visualized utilizing the GRASP system in naïve, single and experienced male flies. The small panels are presented as a red scale to show the GFP fluorescence marked by threshold function of ImageJ. See the MATERIALS AND METHODS for a detailed description of the fluorescence intensity analysis used in this study. (S) GRASP assay for *SIFa^2A-lexA^* and *GAL4^24F06^* in AL region of naïve (top panels), single (middle panels) and experienced (bottom panels) male flies. Scale bars represent 50 μm. (T-W) Quantification of relative value for synaptic area. See the MATERIALS AND METHODS for a detailed description of the fluorescence intensity analysis used in this study. (X) Diagram of various form of connectivity between *SIFa* and *SIFaR* in different sociosexual experience in PI, FB, LAL and AL. The legend above provides the interpretation for each graphic. Larger graphics indicate a higher number of synapses formed. The subsequent diagrams are identical. See also Figure S4.

We utilized synaptobrevin-GRASP (GFP Reconstitution Across Synaptic Partners)^59^, a potent activity-dependent marker for synapses *in vivo* for assessing neuronal connectivity, to examine the synaptic changes between SIFa and SIFaR within brain neurons in various social contexts^60^. Therefore, we employed the SIFa^2A^-lexA driver, which specifically labels ventral-posterior SIFa neurons (SIFa^VP^) that only project to the brain^49^. Using GRASP experiments, we determined that SIFa^VP^-SIFaR neurons form synapses in the gall (GA) of the brain (Fig. 3A). GA are made up of a small group of neurons with numerous synapses and relatively few glia around them. They stretch out from the top of the lateral accessory lobe (LAL), adjacent to the ventrolateral protocerebrum and below the spur of the mushroom body (MB)^61^. Since GA are linked to the EB and protocerebral bridge (PB), they may facilitate learning and memory (Supplementary information, Fig. S4A)^62–64^. We used the “Virtual Fly Brain (VFB)” platform, an interactive tool for investigating neuron connectivity, to confirm the neuronal connectivity of the SIFa neuron with four other neurons, namely GA, FB, and AL (Fig. 3B and supplementary information, Fig. S4B)^65^.

Males reared in groups had stronger connections between SIFa and SIFaR neurons in CB, PI, GA and fan-shaped body (FB) than males reared in isolation (compare naïve and single in Fig. 3C, D; Supplementary information, Fig. 3G, H; Fig. 3K, L; Fig. 3O, P). In addition, the strength of the connections in CB, PI, GA, and FB was reduced in males with sexual experience compared to males without sexual experience (compare nave and exp. in Fig. 3E, F; Fig. 3I, J; Fig. 3M, N; Fig. 3Q, R). In addition to the area of synapses, the number of synapses decreased when males were socially isolated (Supplementary information, Fig. S4C, D) and sexually experienced (Supplementary information, Fig. S4G, H). However, the intensity and size of synapses in different conditions remained constant (Supplementary information, Fig. S4E, F; Fig. S4I, J), indicating that the changes of SIFa-SIFaR synapses by sociosexual experiences are authentic. In contrast to synapses between SIFa and SIFaR formed in PI, GA, and FB, synapses between SIFa and SIFaR neurons in AL remained constant or did not vary significantly across sociosexual conditions (Fig. 3S-W). In VNC, no synapses were found between *SIFa^2A^-lexA* and *GAL4^24F06^* (Supplementary information, Fig. S4K). All of these findings indicate that sociosexual experience may have a substantial effect on the connectivity between SIFa and SIFaR neurons (Fig. 3X).

### Social Context-Induced Synaptic Reorganization Precedes Calcium-Dependent Activity Modifications in SIFaR Neurons and Generate Feed-forward Enhancement

Given the strong expression of SIFaR^2A^ cells in the AG of VNC, we employed *GAL4^SIFa.PT^*, a GAL4 driver line that project to both AG and the brain area^49^, to examine the synaptic connections between SIFa and SIFaR neurons in VNC across various social contexts. Social isolation and sexual experiences led to a decrease in the overall synaptic connections between SIFa and SIFaR in the brain and VNC. This reduction was particularly observed in the decline of OL when socially isolated and AG when both socially isolated and sexually experienced (Fig. 4A-I; Supplementary information, Fig. S5A, B). These findings indicate that social isolation leads to a decrease in the SIFa-SIFaR synapses in the visual pathways (OL), as well as a significant decrease in the SIFa-SIFaR synapses in the AG of VNC when males experience social isolation or sexual interactions (Fig. 4J).

**Figure 4.**
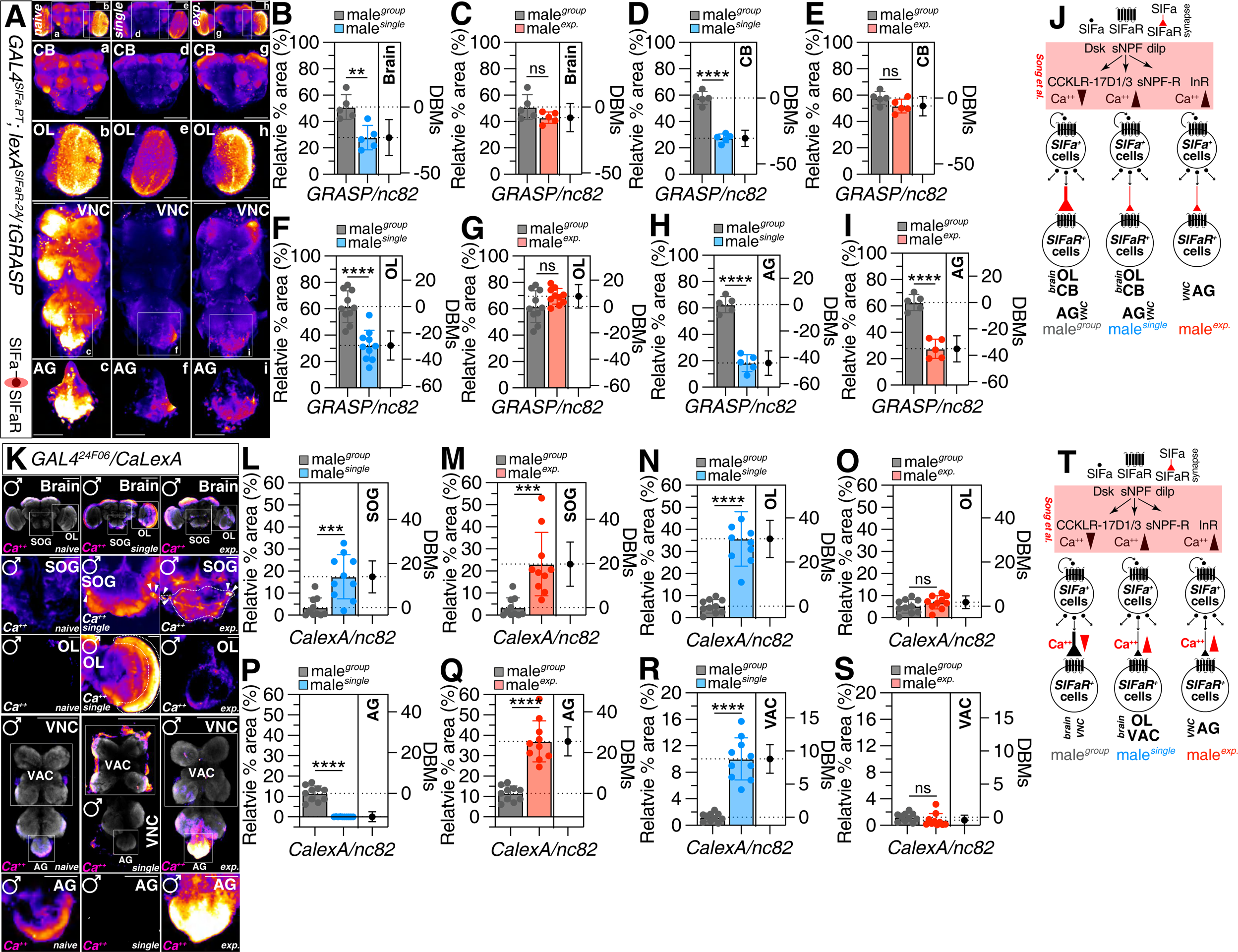
Social context modulates both the formation of synapses between *GAL4^SIFa.PT^* and *lexA^SIFaR-2A^* neurons and the calcium-dependent activity of *SIFaR*. (A) tGRASP assay for *GAL4^SIFa.PT^* and *lexA^SIFaR-2A^* in whole brain, CB, OL, VNC and abdominal ganglion (AG) region of male flies. Male flies expressing *GAL4^SIFa.PT^*, *lexA^SIFaR-2A^* and *lexAop-2-post-t-GRASP, UAS-pre-t-GRASP* were dissected after 5 days of growth. Synaptic transmission occurs from *GAL4^SIFa.PT^* to *lexA^SIFaR-2A^.* GFP is pseudo-colored as “red hot”. Scale bars represent 100 μm upper panels and 50 μm in AG panels. (B-I) Quantification of relative value for synaptic area. See the MATERIALS AND METHODS for a detailed description of the fluorescence intensity analysis used in this study. (J) Schematic representation of the variations in SIFa-SIFaR signaling across distinct regions of the central nervous system (CNS) of male *Drosophila melanogaster* under diverse social contexts. (K) Different levels of neural activity of the brain as revealed by the CaLexA system in naïve, single and experienced flies. Male flies expressing *GAL4^24F06^* along with *LexAop-CD2-GFP, UAS-mLexA-VP16-NFAT and LexAop-CD8-GFP-A2-CD8-GFP* were dissected after 5 days of growth (mated male flies had 1-day of sexual experience with virgin females). The dissected brains were then immunostained with anti-GFP (green) and anti-nc82 (blue). GFP is pseudo-colored as “red hot”. Boxes indicate the magnified regions of interest presented in the bottom panels. Scale bars represent 100 μm in brain, VNC and SOG panels, 25 μm in OL panels, and 50 μm in AG panels. (L-S) Quantification of relative value for GFP fluorescence. See the MATERIALS AND METHODS for a detailed description of the fluorescence intensity analysis used in this study. (T) Illustration depicting the variations in activity of SIFaR^+^ neurons within various regions of the CNS across different social contexts in male *Drosophila*. The red arrow denotes the calcium ion concentration, with subsequent diagrams following the same representation. See also Figure S5.

Next, we investigated the calcium response properties of *GAL4^24F06^* neurons in different social experiences of fly. We analyzed neuronal activity using a CaLexA-based transcriptional reporter system in which sustained neural activity drives GFP expression^66^. We found that the levels of CaLexA signals in both the brain and VNC were significantly elevated in both socially isolated and sexually experienced males when compared to group reared males (Supplementary information, Fig. S5E-H). CaLexA signals originating from *GAL4^24F06^*-positive SOG neurons were found to be at higher levels in males reared singly and with sexual experience than in males reared in groups (Fig. 4L, M). In comparison to the SOG region, the calcium signals observed in SIFaR^24F06^-positive OL neurons exhibited an increase solely in the singly reared condition, while no significant difference was observed in the sexually experienced condition when compared to the group reared condition (Fig. 4N, O).

The CaLexA signals originating from the AG region in the VNC showed a decrease when males were subjected to social isolation (Fig. 4P). Conversely, these signals exhibited an increase when males had prior sexual experience (Fig. 4Q). In contrast to the calcium signal changes of AG, the CaLexA signals originating from the VAC region exhibited an increase when males were socially isolated, but no significant differences were observed when sexually experienced (Fig. 4R, S). These findings suggest a strong link between the synaptic plasticity of SIFa-SIFaR and calcium activity in GAL4^24F06^-positive neurons in males, in response to different social context^49^ (Fig. 4T).

Additionally, SIFa neurons express SIFaR, which enables them to produce SMD behavior^49^. In order to assess the influence of significant alterations in calcium activity that occur precede SIFa-SIFaR synaptic changes, we examined the SIFaR-to-SIFa synaptic changes using tGRASP. No synapses along the SIFaR-to-SIFa direction were observed in the VNC (Supplementary information, Fig. S5C, D and I). Significant synaptic plasticity has been observed between SIFaR and SIFa neurons in the PI, PRW, and OL regions of the brain. In general, social isolation led to a decrease in SIFaR-SIFa synapses, while sexual experiences resulted in an increase (Supplementary information, Fig. S5J, K). Additionally, synapses around the PI region also showed an increase (Supplementary information, Fig. S5L, M). Social isolation and sexual experiences significantly reduce the synapses near the prow (PRW) region, where it contains the circuitry underlying feeding behavior and is involved in numerous other aspects of sensory processing and motor control^67^. The synapses between SIFaR and SIFa in OL exhibited a decrease solely in males who were socially isolated, rather than in those who had sexual experiences (Supplementary information, Fig. S5P, Q). Therefore, it can be demonstrated that the synaptic modifications of SIFa-to-SIFaR take place when there are changes in the calcium activity of SIFaR neurons, resulting in changes in SIFaR-to-SIFa synapses (Supplementary information, Fig. S5R). The evidence indicates that neuronal circuits involving SIFa-to-SIFaR-to-SIFa utilize feed-forward augmentation to modify the internal states of the CNS in response to different social contexts (Supplementary information, Fig. S5S, T).

### Corazonin relays the SIFa signaling pathway to SIFaR and CrzR-expressing neurons and glia for the generation of distinct interval timing behavior

The neuropeptide Corazonin (Crz) has been identified as a participant in the regulation of male *Drosophila* mating duration within the AG of the VNC. The male AG’s four neurons that express Crz also express fruitless (fru), and their silencing results in reduced sperm transfer and extended mating duration^68^. Neurons expressing Crz exhibit robust synaptic connections with SIFaR^24F06^ neurons located in the PRW region of the SOG in the brain (panels of Brain and SOG in Fig. 5A). It is noteworthy that approximately 65% and 90% of neuronal signals that tested positive for SIFaR^24F06^ also exhibited co-expression of Crz-GAL4 in the brain and outer region of the OL, respectively (Supplementary information, Fig. S6A, B; Fig. 5B). However, the percentage of neuronal signals exhibiting co-expression in the CB or SOG region was only 30% (Fig. 5C and supplementary information, Fig. S6E). The modulation of interval timing in LMD is primarily facilitated through the visual pathway^36,37,46^. It is plausible that the expression of Crz in SIFaR^24F06^ neurons located in the OL may play a crucial role in the manifestation of LMD behavior. In comparison to the brain, a smaller proportion of SIFaR^24F06^ signals exhibited overlap with *Crz-GAL4* in VNC, amounting to only 10% (Supplementary information, Fig. S6C, D). However, this percentage increased to 30% in the AG region (Fig. 5D, E). The signals of SIFaR^24F06^ located outside of the AG region in VNC exhibited minimal overlap with Crz signals (Fig. 5D).

**Figure 5.**
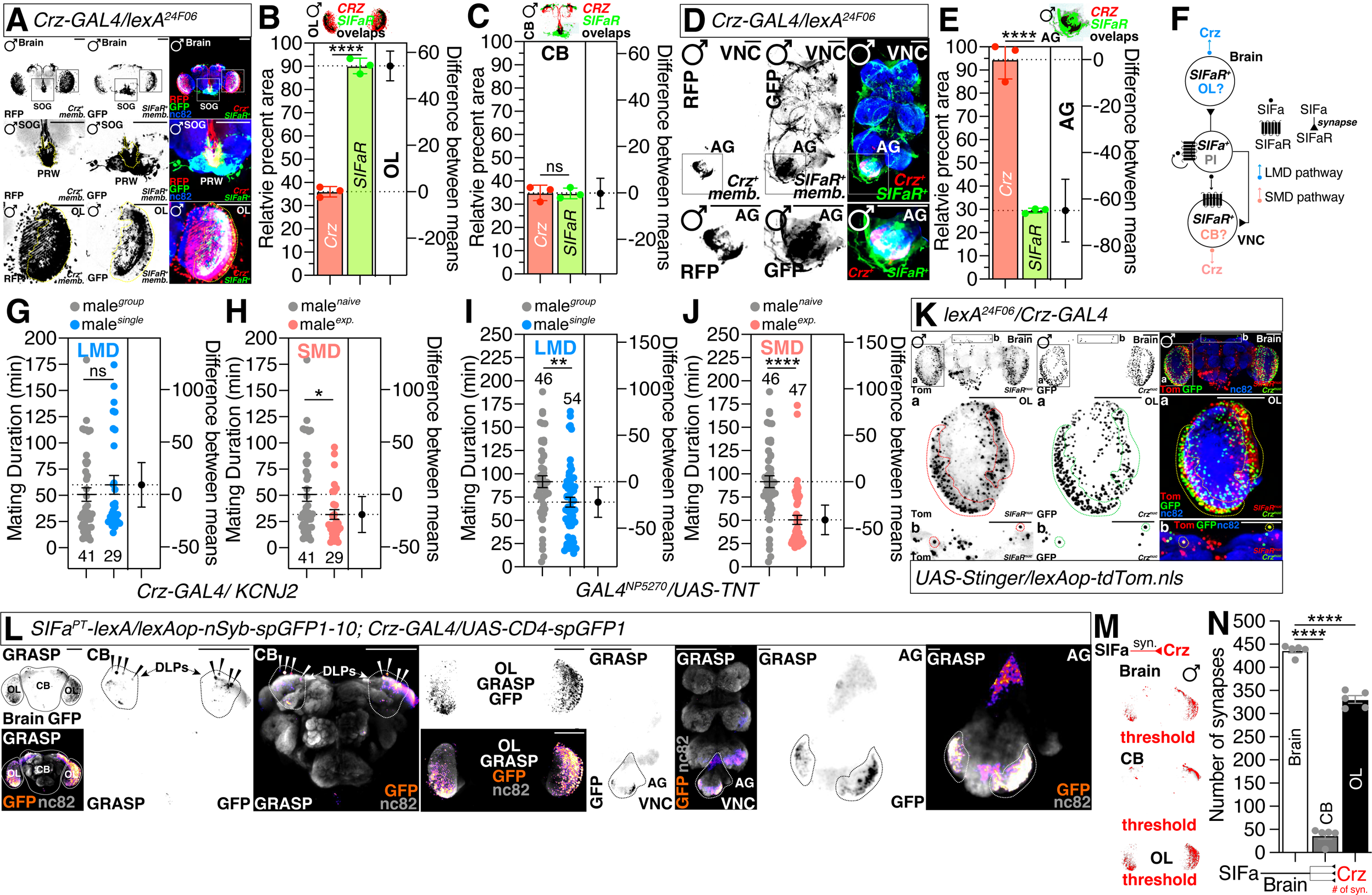
*Crz* modulate LMD and SMD behaviors through SIFa-SIFaR signaling. (A) Male flies brain expressing *Crz-GAL4* and *lexA^24F06^* drivers together with *UAS-mCD8RFP* and *lexAop-mCD8GFP*. Boxes indicate the magnified regions of interest presented in the bottom panels. Scale bars represent 100 μm. (B-C) Colocalization analysis of GFP and RFP staining, normalized to total GFP and RFP areas. See the MATERIALS AND METHODS for a detailed description of the colocalization analysis used in this study. (D) Male flies VNC expressing *Crz-GAL4* and *lexA^24F06^* drivers together with *UAS-mCD8RFP* and *lexAop-mCD8GFP*. Scale bars represent 50 μm. (E) Colocalization analysis of GFP and RFP staining, normalized to total GFP and RFP areas. See the MATERIALS AND METHODS for a detailed description of the colocalization analysis used in this study. (F) Illustration depicting the mechanism by which Crz modulates LMD and SMD via SIFa-SIFaR signaling pathway. (G-H) LMD and SMD assays for *Crz-GAL4* mediated electrical silencing *via UAS-KCNJ2*. (I-J) LMD and SMD assays for *GAL4^NP5270^* mediated inactivation of synaptic transmission *via UAS-TNT*. (K) Male flies brain expressing *Crz-GAL4* and *lexA^24F06^* drivers together with *UAS-Stinger* and *lexAop-tdTomato.nls*. Boxes indicate the magnified regions of interest presented in the bottom panels; dashed circles indicate overlapped cells. Scale bars represent 100 μm. (L) GRASP assay for *SIFa^PT^-lexA* and *Crz-GAL4* in whole brain, CB, OL, VNC and AG region of male flies. GFP is pseudo-colored as “Magenta hot”. Synaptic transmission occurs from *SIFa^PT^-lexA* to *Crz-GAL4.* Scale bars represent 100 μm in left panels and 10 μm in AG panels. (M) The GFP fluorescence (green) in male fly brain was processed using ImageJ software, where a threshold function was applied to distinguish fluorescence from the background. (N) Quantification of synapses formed between *SIFa^PT^-lexA* and *Crz-GAL4* in brain, CB and OL. See the MATERIALS AND METHODS for a detailed description of the particle analysis used in this study. See also Figure S6.

We have previously determined that Crz expression in SIFaR^24F06^ cells is required for both LMD and SMD (Supplementary information, Fig. S6F, G), whereas SIFaR expression in Crz neurons is only required for SMD and not LMD (Supplementary information, Fig. S6H, I)^46^. It has been reported that KCNJ2, which functions by inwardly rectifying the K^+^ channel to impede Crz neuronal activity, can substantially extend the duration of mating^68^. Previous research has documented a significant augmentation in the mating duration exhibited by these males. However, this genetic manipulation only disrupted LMD and not SMD (Fig. 5G, H).

A limited number of dsx-positive neurons situated in AG have been recognized, which are composed of GABAergic and dopaminergic neurons^69^ named as ‘Copulation Decision Neuron (CDN)’^70–72^. CDN function in opposition to regulate copulation duration by modulating the behavioral state of the male fly over a period of time^68,69^. To examine the function of CDN in LMD/SMD behaviors, the *GAL4^NP5270^* neurons were inactivated by expressing tetanus toxin light chain (TeTxLC or TNT). As per prior documentation, there was a notable increase in the duration of mating among these males. However, both LMD and SMD were observed to be normal (Fig. 5I, J). It’s interesting to note that only SMD was affected by the *GAL4^NP5270^*–mediated knockdown of SIFaR, indicating that SIFaR signaling in these cells is essential for controlling SMD behavior rather than LMD behavior (Supplementary information, Fig. S6J, K). These data indicate that the suppression of CDN does not elicit any impact on either LMD or SMD behavior. However, the knockdown of SIFaR in these cells exclusively influences SMD behavior. It is worth noting that the expression of SIFaR in Crz cells is only essential for the SMD behavior (Supplementary information, Fig. S6H, I). These findings collectively suggest that the signaling pathways of CDN and Crz through SIFaR play a more significant role in the generation of SMD behavior compared to neurotransmitter release or the electrical activity of CDN and Crz neurons (Supplementary information, Fig. S6L).

By employing the genetic intersectional method, we have determined that approximately 100 neurons in the OL and 2 neurons in dorsal lateral peptidergic neurons (DLPs) express Crz in conjunction with SIFaR^24F06^ neurons (Fig. 5K)^73^. However, the Crz-positive AG neurons in the VNC are not associated with the SIFaR^24F06^ driver (Supplementary information, Fig. S6M), indicating that only Crz neurons in the brain are positive for SIFaR^24F06^.

There is a strong connection between SIFa neurons and Crz neurons in the OL and DLPs in the brain, as well as in the lateral region of AG in the VNC (Fig. 5L). The number and area of synapses between SIFa and Crz exhibit a significantly greater magnitude in OL compared to CB (Fig. 5M, 5N; Supplementary information, Fig. S6N). Nevertheless, the size of these synapses is similar (Supplementary information, Fig. S6O). The synapses between SIFa and Crz exhibit greater quantity, extent, and dimensions in AG compared to other VNC regions (Supplementary information, Fig. S6P-S). These findings indicate that SIFa neurons establish intact connections with Crz neurons in both the brain and VNC. However, the transmission of signals through SIFaR in Crz-expressing neurons is limited to the brain.

To elucidate the response of Crz neurons to the activity of SIFa neurons, we conducted live calcium (Ca^2+^) imaging in the AG region of the VNC, where Crz neurons are situated (Fig. 6A). We monitored Ca^2+^ dynamics in these neurons using two-photon microscopy with a genetically encoded calcium indicator, GCaMP7s, under the control of the *Crz-GAL4* driver. To selectively activate SIFa neurons, we employed an optogenetic approach using the red-light-sensitive channelrhodopsin (CsChrimson), which was expressed via the *lexA^SIFa.PT^* driver (Fig. 6B). Upon optogenetic stimulation of SIFa neurons, we observed a significant increase in the activity of Crz, evidenced by a sustained elevation in intracellular Ca^2+^ levels that persisted in a high level before gradually declining to baseline levels. This increase was in contrast to the control group (without all-trans retinal, ATR) (Fig. 6C-E).

**Figure 6.**
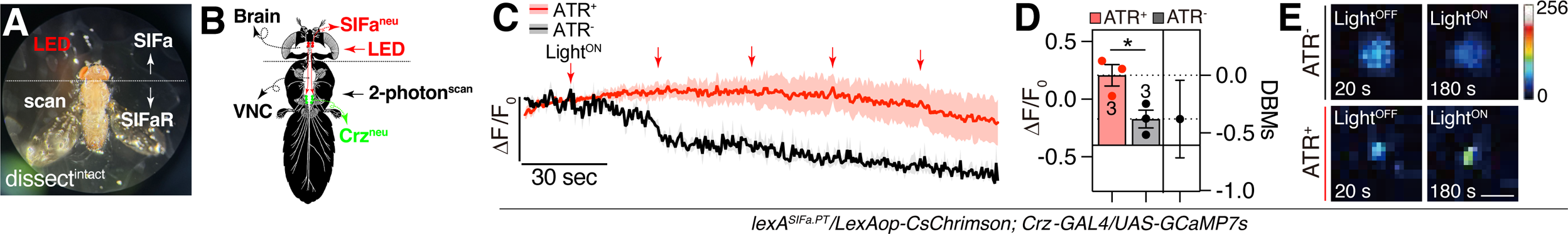
Crz neurons located in the AG are activated by SIFa neuronal activity and play a role in the neural computation of interval timing. (A) An illustrative depiction of a fly immobilized on a plastic plate using UV adhesive on its back. LED light stimulation targeting SIFa neurons in the brain region demarcated by the dashed line. Scanning for neuronal activity was conducted in the area below the dashed line. (B) Schematic diagram depicting the precise location of the two-photon scanning procedure. Green labels indicate the nucleus of Crz neurons located in AG region. Red labels indicate the nucleus of SIFa neurons located in PI region. (C-E) Crz neurons respond to SIFa neurons’ activity. Fluorescence changes (Δ*F*/*F*_0_) of GCaMP7s in Crz neurons after optogenetic stimulation of SIFa neurons. LED light were fired 2 s after 30 s of dark. *N* = 3 in each group. See the MATERIALS AND METHODS for a detailed description of the calcium imaging used in this study.

These findings confirm that Crz neurons are activated in response to SIFa neuronal activity, reinforcing their role as postsynaptic effectors in the neural circuitry governed by SIFa neurons. Moreover, these results provide empirical support for the hypothesis that SIFa-SIFaR/Crz-CrzR long-range neuropeptide relay underlies the neuronal activity-based measurement of interval timing.

Next, to determine how Crz neurons can modulate LMD and SMD behaviors in various brain and VNC regions, we examined the calcium response properties of Crz-expressing neurons in various internal states (Fig. 7A and 7D). An increase in CaLexA signals originating from the OL area in the brain was observed when males were exposed to social isolation (Fig. 7B). In contrast, the above-mentioned signals showed a decline in cases where males had previous sexual experiences (Fig. 7C). The collective CaLexA signals observed across the entire brain and CB exhibit a similar pattern to that of the OL (Supplementary information, Fig. S7A-D). This suggests that the calcium levels of Crz neurons in the majority of brain regions increase during social isolation and decrease following sexual experience in males. In contrast to the calcium signal changes of brain region, the CaLexA signals originating from the AG (Fig. 7E-F) and VNC (Supplementary information, Fig.S7E-F) exhibited an increase when males were sexually experienced, but no significant differences were observed when socially isolated. These data indicate that there is a significant change in calcium activity among Crz-positive OL and AG cells in response to different social experiences in males.

**Figure 7.**
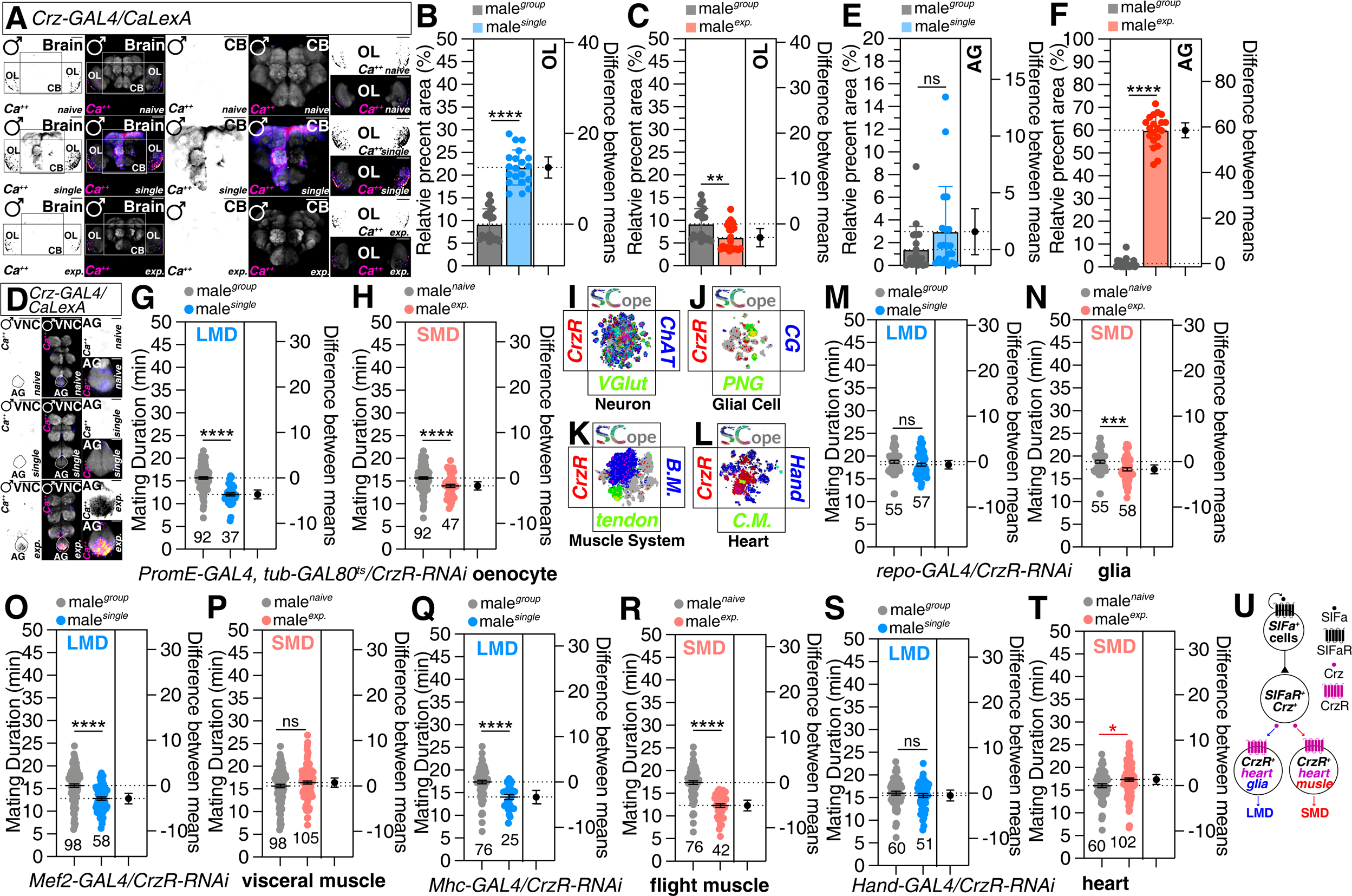
Social context modulates the calcium-dependent activity of *Crz* and the expression of CrzR in non-neuronal tissue is associated with LMD and/or SMD. (A) Different levels of neural activity of the brain as revealed by the CaLexA system in male flies. GFP is pseudo-colored as “red hot”. Boxes indicate the magnified regions of interest presented in the bottom panels. Scale bars represent 100 μm. (B-C) Quantification of GFP fluorescence. See the MATERIALS AND METHODS for a detailed description of the fluorescence intensity analysis used in this study. (D) Different levels of neural activity of the VNC as revealed by the CaLexA system in male flies. Scale bars represent 100 μm in VNC panels and 25 μm in AG panels. (E-F) Quantification of GFP fluorescence. See the MATERIALS AND METHODS for a detailed description of the fluorescence intensity analysis used in this study. (G-H) LMD and SMD assays for *PromE-GAL4* mediated knockdown of SIFaR *via SIFaR-RNAi*. (I-L) Single-cell RNA sequencing (SCOPE scRNA-seq) datasets reveal cell clusters colored by expression of *CrzR* (red), *ChaT/VGlut* (blue/green) in neurons (I), *CG* (cortex glia)*/PNG* (perineurial glia) (blue/green) in glial cells (J), *B.M* (body muscle)*/tendon* (blue/green) in muscle systems (K), and *Hand/C.M* (blue/green) in heart clusters (L). (M-T) LMD and SMD assays for *repo-GAL4* (glia), *Mef2-GAL4* (visceral muscle), *Mhc-GAL4* (flight muscle) and *Hand-GAL4* (heart) mediated knockdown of SIFaR *via UAS-SIFaR-RNAi.* Red asterisk denotes a significant effect of the experiment on SMD, leading to prolonged mating duration in experienced vs. naïve male flies. (U) Diagram depicting the necessity of non-neuronal populations for the SIFa-SIFaR/Crz-CrzR neuropeptide signaling pathway in regulating LMD and SMD. See also Figure S7.

### The expression of the Crz receptor (CrzR) in non-neuronal tissue is associated with the display of different interval timing behaviors

The impact of Crz is facilitated through the corazonin receptor (CrzR) and projection neurons that express CrzR, which are responsible for innervating the male reproductive organs and promoting the timing of sperm and seminal fluid transfer (SSFT) through the release of 5-HT^68^. The CrzR gene is expressed in both salivary glands and adipocytes of the fat body and is involved in regulating resistance to various stressors such as starvation, desiccation, oxidative stress, feeding, and ethanol-related behaviors^74–76^. The majority of CrzR expression can be found in oenocytes (Supplementary information, Fig. S7G), and adult oenocytes (Supplementary information, Fig. S7H) have an enrichment of CrzR, according to the analysis of the fly Scope scRNA sequencing dataset^77^. However, the suppression of CrzR in oenocytes did not have any impact on both LMD and SMD behaviors in adults (Fig. 7G, H), indicating that oenocyte CrzR is not necessary for interval timing behaviors.

Remarkably, the pan-neuronal suppression of CrzR exhibits no discernible impact on either LMD or SMD (Supplementary information, Fig. S7I, J; Fig. 7I). Interestingly, our findings indicate that the downregulation of CrzR in the glial population resulted in the disruption of LMD rather than SMD (Fig. 7M, N). The expression of CrzR in glial cells was predominantly observed in the perineurial glia population (Fig. 7J). Mef2-GAL4-mediated knockdown of CrzR in muscle only affects SMD; in contrast, Mhc-GAL4-mediated knockdown had no effect on SMD, indicating that CrzR expression in visceral muscle, not flight muscle, is necessary for SMD behavior (Fig. 7O-R). It is noteworthy that the majority of CrzR expression within the muscle system is observed in tendon cells and body muscle (Fig. 7K). The RNAi knockdown of CrzR in adult fat body (Supplementary information, Fig. S7K, L), gut enterocyte (Fig. S7M-N), or salivary gland (Fig. S7O, P) had no effect on both LMD and SMD behaviors. The knockdown of CrzR in the heart via Hand-GAL4 disrupts both LMD and SMD behaviors (Fig. 7S, T). This suggests that the expression of CrzR in the cardiomyocyte of the heart (Fig. 7L) might play a crucial role in modulating interval timing behaviors. All these data indicate that the relay of SIFa-SIFaR/Crz-CrzR neuropeptide signaling necessitates the presence of non-neuronal population in order to regulate interval timing behaviors (Fig. 7U).

## DISCUSSION

Here we delved into the mechanisms by which the neuropeptide SIFa and its receptor SIFaR regulate two distinct interval timing behaviors in male *Drosophila melanogaster*: Longer-Mating-Duration (LMD) and Shorter-Mating-Duration (SMD). Utilizing RNAi and genetic rescue techniques, we identified that the expression of SIFaR in specific neuronal populations is crucial for sustaining LMD and SMD (Fig. 1 and 2). We further explored the synaptic connections between SIFa and SIFaR neurons, uncovering that social context and sexual experience can lead to synaptic reorganization, thereby influencing the internal states of the central nervous system (Fig. 3 and 4). Our study provides evidence that neuropeptide relay pathways involving SIFa-SIFaR/Crz-CrzR are integral in generating these distinct interval timing behaviors. Additionally, the activity of Crz neurons, which is modulated by SIFa neurons, plays a role in the neuronal mechanisms underlying the measurement of interval timing (Fig. 5-7). These insights contribute to a broader understanding of the neural circuitry that underlies interval timing, with potential implications for a range of behavioral adaptations modulated by neuropeptidergic system.

Neuropeptides encompass a broad and heterogeneous group of signaling molecules that are synthesized by various cellular entities. As neuromodulators, co-transmitters, or circulating hormones, various neuropeptides have pleiotropic effects. However, there is a lack of understanding of the mechanisms by which neuropeptides can exhibit pleiotropic roles in context-dependent behaviors. The fact that numerous neuropeptides exhibit pleiotropic effects does not prevent us from gaining a comprehensive grasp of their functions at the organismal level. The autonomous functioning of peptidergic neurons is infrequent, as they mostly incorporate the modulatory influences of other peptidergic systems. Nevertheless, the process of achieving balance in this integration remains poorly understood^3,73^.

SIFa-producing neurons are composed of a distinct quartet of neurons that exhibit extensive arborization across the majority of brain regions and AG. Therefore, the SIFa system is regarded as a typical neuropeptidergic system due to its ability to incorporate internal signals that are dependent on the state or context into the orchestrating peptidergic system. SIFa neurons integrate multiple inputs, which may contain encoded external and internal cues, as well as fundamental conditions like metabolic status, across numerous circuits. These particular large peptidergic neurons have the capability to function as components of circuits that assess internal states and sensory inputs, thereby establishing transitions between conflicting behaviors such as mating and aggression or feeding and sleep^4,7,49^.

An obstacle in understanding the complex role of neuropeptides is the scarcity of information regarding the cellular arrangement of neuropeptide receptors, which directly influence neuromodulatory circuits. Except for a few cases, we lack knowledge regarding the precise location of the receptor protein. In this study, we demonstrated the ability of a certain subset of neurons expressing SIFaR to selectively regulate two unique interval timing behaviors. Additionally, our findings demonstrate that the neuropeptide relay mediated by SIFa-SIFaR/Crz-CrzR has the potential to serve as a viable model for modulating context-dependent behavior through alterations in synaptic plasticity and calcium activity. Additionally, our study demonstrated that CrzR-expressing cells in non-neuronal populations play a distinct role in interval timing behaviors. This finding implies that the neuropeptide relay mechanism is not solely dependent on communication among neurons, but also encompasses communication between neurons and non-neuronal tissues.

The presence of multiple different neurons that regulate the duration of mating in AG has been reported. According to Taylor et al^68^, the use of KCNJ2 to block fru-positive Crz expressing AG neurons results in an increase in mating duration up to 100 minutes. Furthermore, the use of TNT to suppress dsx-positive CDN neurons in AG has been found to extend the mating duration by up to 100 minutes^69,71^. Our findings demonstrate that cells expressing SIFaR in AG play a crucial role in regulating LMD and SMD behaviors. Due to the absence of fru- or dsx-positive neurons in SIFaR^24F06^ cells, we classify these cell types as separate from previously documented Crz or CDN neurons. Notably, we observed that SMD behavior is unaffected in both scenarios when Crz and CDN neurons significantly prolong the mating duration (Fig. 5G-J). This observation suggests that the modulation of mating duration by Crz and CDN can still be influenced by neuropeptide relay signals from the PI of SIFa to AG of SIFaR. This finding further demonstrates that the transmission of neuropeptides through SIFa can indeed induce behavior that is context-dependent, irrespective of the downstream neuronal circuits involved.

Neuropeptide relays are fundamental for survival and goal-oriented behaviors. One apparent case is the balance between appetite and satiety via orexigenic and anorexigenic neuronal circuits. It has been known that neuropeptides and hormones are important neural substrates for modulating appetite and satiety control^78,79^. SIFa also has been known as a central modulator in parallel inhibition between orexigenic and anorexigenic pathways^80^. Further investigations on SIFa-mediated neuropeptide relay will provide a foundational understanding of how neuropeptides and their receptors modulate neuronal circuits.

## MATERIALS AND METHODS

### Fly Stocks and Husbandry

*Drosophila melanogaster* were raised on cornmeal-yeast medium at similar densities to yield adults with similar body sizes. Flies were kept in 12 h light: 12 h dark cycles (LD) at 25 (ZT 0 is the beginning of the light phase, ZT12 beginning of the dark phase) except for some experimental manipulation (experiments with the flies carrying tub-GAL80^ts^). Wild-type flies were Canton-S.

Following lines used in this study, *Canton-S* (#64349), *Df(1)Exel6234* (#7708), *Crz-GAL4* (#51977), *elav^c155^; UAS-Dicer* (#25750), *lexAop-CD8GFP; UAS-mLexA-VP16-NFAT, lexAop-rCD2-GFP* (#66542), *lexAop-nSyb-spGFP1-10, UAS-CD4-spGFP11* (#64315), *UAS-post-t-GRASP, lexAop2-pre-t-GRASP* (#79039), *UAS-pre-t-GRASP, lexAop2-post-t-GRASP* (#79040), *PromE-GAL4* (#65405), *repo-GAL4* (#7415), *Sgs3-GAL4* (#6870), *SIFaR^B322^* (#16202), *GAL4^23G06^* (#49041), *GAL4^24A12^* (#49061), *GAL4^24F06^* (#49087), *GAL4^57F10^* (#46391), *lexA^24F06^* (#52695), *SIFaR-RNAi^JF01849^* (#25831), *SIFaR-RNAi^HMS00299^* (#34947), *tub-GAL80^ts^* (#7108), *UAS-CD4tdGFP*(#35839), *UAS-RedStinger* (#8546), *CrzR-RNAi^JF02042^* (#26017), *UAS-Denmark, UAS-syt.eGFP* (#33065), *UAS-KCNJ2* (#6596), *UAS-mCD8RFP, lexAop-mCD8GFP* (#32229), *UAS-GCaMP7s* (#80905), *lexAop-CsChrimson* (#55138), *UAS-TNT* (#28838), *Mhc-GAL4* (#55133), *Hand-GAL4* (#48396), *Mef2-GAL4* (#27390), *3.1Lsp2-GAL4* (#84285), and *Myo31DF-GAL4* (#84307) were obtained from the Bloomington *Drosophila* Stock Center at Indiana University. The following lines, *SIFaR-RNAi^v1783^*(#1783), and *Crz-RNAi^v30670^* (#30670) were obtained from the Vienna *Drosophila* Resource Center. *GAL4^NP5270^* was obtained from KYOTO *Drosophila* Stock Center. The following lines, *SIFa-lexA^T2A^* (#FBA00116), *SIFa-GAL4^T2A^* (#FBA00103), *SIFaR-GAL4^T2A^* (#FBF00102), and *SIFaR-lexA^T2A^* (#FBF00086) were obtained from Qidong Fungene Biotechnology in China. The *GAL4^SIFa.PT^* was a gift from Jan A. Veenstra. The *UAS-SIFaR* was a gift from Young-Joon Kim in GIST. To reduce the variation from genetic background, all flies were backcrossed for at least 3 generations to CS strain. All mutants and transgenic lines used here have been described previously.

### Mating duration assay

The mating duration assay in this study has been reported^36,37,48^. To enhance the efficiency of the mating duration assay, we utilized the *Df(1)Exel6234* (DF here after) genetic modified fly line in this study, which harbors a deletion of a specific genomic region that includes the sex peptide receptor (SPR)^81,82^. Previous studies have demonstrated that virgin females of this line exhibit increased receptivity to males^82^. We conducted a comparative analysis between the virgin females of this line and the CS virgin females and found that both groups induced SMD. Consequently, we have elected to employ virgin females from this modified line in all subsequent studies. For naïve males, 40 males from the same strain were placed into a vial with food for 5 days. For single reared males, males of the same strain were collected individually and placed into vials with food for 5 days. For experienced males, 40 males from the same strain were placed into a vial with food for 4 days then 80 DF virgin females were introduced into vials for last 1 day before assay. 40 DF virgin females were collected from bottles and placed into a vial for 5 days. These females provide both sexually experienced partners and mating partners for mating duration assays. At the fifth day after eclosion, males of the appropriate strain and DF virgin females were mildly anaesthetized by CO_2_. After placing a single female in to the mating chamber, we inserted a transparent film then placed a single male to the other side of the film in each chamber. After allowing for 1 h of recovery in the mating chamber in 25 incubator, we removed the transparent film and recorded the mating activities. Only those males that succeeded to mate within 1 h were included for analyses. Initiation and completion of copulation were recorded with an accuracy of 10 sec, and total mating duration was calculated for each couple. All assays were performed from noon to 4pm. We conducted blinded studies for every test.

### Generation of transgenic flies

To generate the *SIFa^PT^-lexA* driver, the putative promotor sequence of the gene was amplified by PCR using wild-type genomic DNA as a template with the following primers GCCAATTGGCTGAATCTCCTGACCCTCA and GCAGATCTCTTGCAGTTTTCGGTGAGC as mentioned before^83^. The amplified DNA fragment (1482 base pairs located immediately upstream of the *SIFa* coding sequence) was inserted into the E2 Enhancer-lexA vector. This vector, supplied by Qidong Fungene Biotechnology Co., Ltd. (http://www.fungene.tech/), is a derivative of the pBPLexA::p65Uw vector (available at https://www.addgene.org/26231). The insertion was achieved by digesting the fragment and the vector with EcoRI and XbaI restriction enzymes to create compatible sticky ends. The genetic construct was inserted into the *attp2* site on chromosome III to generate transgenic flies using established techniques, a service conducted by Qidong Fungene Biotechnology Co., Ltd.

### Immunostaining

The dissection and immunostaining protocols for the experiments are described elsewhere^37^. After 5 days of eclosion, the *Drosophila* brain will be taken from adult flies and fixed in 4% formaldehyde at room temperature for 30 minutes. The sample will be washed three times (5 minutes each) in 1% PBT and then blocked in 5% normal goat serum for 30 minutes. The sample will next be incubated overnight at 4 with primary antibodies in 1% PBT, followed by the addition of fluorophore-conjugated secondary antibodies for one hour at room temperature. The brain will be mounted on plates with an antifade mounting solution (Solarbio) for imaging purposes. Samples were imaged with Zeiss LSM880. Antibodies were used at the following dilutions: Chicken anti-GFP (1:500, Invitrogen), mouse anti-nc82 (1:50, DSHB), rabbit anti-DsRed (1:500, Rockland Immunochemicals), Alexa-488 donkey anti-chicken (1:200, Jackson ImmunoResearch), Alexa-555 goat anti-rabbit (1:200, Invitrogen), Alexa-647 goat anti-mouse (1:200, Jackson ImmunoResearch).

### Quantitative analysis of fluorescence intensity

To quantify the calcium level and synaptic intensity in microscopic images, we introduced ImageJ software^84^. We initially employed ImageJ’s ‘Measure’ feature to calculate average pixel intensity across the entire image or in user-specified sections, and the ‘Plot Profile’ feature to create intensity profiles across areas. To maximize precision, we converted color images to grayscale before analysis. Thresholding methods were also utilized to produce binary images that accurately outlined areas of interest, with pixel intensities of 255 (white) assigned to regions of interest and 0 (black) to the background. Intensity values from the binary image were then transferred to the corresponding locations in the original grayscale image to obtain precise intensity measurements for each object. The ‘Display Results’ feature provided comprehensive data for each object, including average intensity, size, and other relevant statistics. To normalize for fluorescence differences between ROIs, GFP fluorescence for CaLexA and tGRASP was normalized to nc82. All specimens were imaged under identical conditions.

### Colocalization analysis

To perform colocalization analysis of multi-color fluorescence microscopy images in this study, we employed ImageJ software^85^. Initially, merge the individual channels of the fluorescence image to create a composite view. Following this, access the ‘Image’ menu, select ‘Type’, and then choose ‘RGB Color’ to ensure the image is displayed in the correct color format. Next, navigate to ‘Image’, then ‘Adjust’, and select ‘Color Threshold’. Adjust the threshold settings to isolate the yellowish pixels indicative of colocalization, and confirm the selection by clicking the ‘Select’ button. Subsequently, proceed to ‘Analyze’, followed by ‘Measure’. This action will open a new window displaying the measurement data for the selected area. Record the ‘Area’ value provided in the measurement window, as this corresponds to the area of colocalization between the two fluorophores. To calculate the percentage of the colocalized area relative to the total area of the fluorophore of interest (e.g., GFP or RFP), readjust the threshold settings to encompass the entire fluorophore area and click ‘Select’ again. Afterward, repeat the measurement process by going to ‘Analyze’ and then ‘Measure’. This will yield the total area value for the respective fluorophore. Finally, to determine the colocalization percentage, divide the area value of the colocalized region by the total area value of the GFP or RFP region. This calculation provides the colocalization efficiency within the specified region. Additionally, we displayed images of fluorescence and overlapping areas in the male fly brain and/or VNC, processed with ImageJ software using a threshold function to differentiate fluorescence from background. All specimens were imaged under identical conditions.

### Particle analysis

To quantitatively measure particle intensity of cell number and synaptic puncta in microscopic images, we applied ImageJ software^86^. Initially, the image is converted to grayscale to reduce complexity and enhance contrast. Subsequently, thresholding techniques are employed to binarize the image, distinguishing particles from the background. This binarization can be achieved through automated thresholding algorithms or manual adjustment to optimize the segmentation. Following binarization, the “Analyze Particles” function is utilized to enumerate and measure the particles based on user-defined criteria, such as size constraints and shape descriptors (e.g., area, perimeter, circularity). The results of these measurements are then available for review in the “Results” window, and the particles can be labeled within the image for visual identification. All specimens were imaged under identical conditions.

### Two-photon Calcium Imaging

Male flies of proper genotype were selected and separated into two groups (with and without ATR) after 5 days of eclosion. The flies were carefully sedated with ice and then placed in an inverted posture on a plastic plate coated with UV glue, which securely held flies in the middle. The plate was filled with *Drosophila* Adult Hemolymph-Like Saline (AHLS) buffer [108 mM NaCl, 5 mM KCl, 4 mM NaHCO_3_, 1 mM NaH_2_PO_4_, 15 mM ribose, mM Hepes (pH 7.5), 300 mosM, CaCl_2_ (2 mM), and MgCl_2_ (8.2 mM)] to ensure the flies’ neurons remained in an active state throughout the experiment^87,88^. Subsequently, the flies were dissected to specifically expose the VNC for detailed examination. Calcium imaging was performed using Zeiss LSM880 microscope with a 20x water immersion objective. GCaMP7 slow (GCaMP7s) signals were recorded with an 880 nm laser and optogenetic stimulation was achieved with a 590 nm LED. LED stimulation paradigm: 30 s off, 2 s on, 30 s off, 2 s on, 30 s off, 2 s on, 30 s off, 2 s on, 30 s off, 2 s on, 30 s off. ROIs were manually selected from the Crz cell body in AG area with ImageJ. Fluorescent change was calculated as *(F*_peak_ *− F_0_)/F_0_*, where *F_0_* was calculated from the average intensity of 10 frames of background-subtracted baseline fluorescence before optogenetic stimulation, and *F*_peak_ corresponds to the highest fluorescence after stimulation.

### Single-nucleus RNA-sequencing analyses

snRNAseq dataset analyzed in this paper is published^89^ and available at the Nextflow pipelines (VSN, https://github.com/vib-singlecell-nf), the availability of raw and processed datasets for users to explore, and the development of a crowd-annotation platform with voting, comments, and references through SCope (https://flycellatlas.org/scope), linked to an online analysis platform in ASAP (https://asap.epfl.ch/fca). Single-cell RNA sequencing (scRNA-seq) data from the *Drosophila melanogaster* were obtained from the Fly Cell Atlas website (https://doi.org/10.1126/science.abk2432). Oenocytes gene expression analysis employed UMI (Unique Molecular Identifier) data extracted from the 10x VSN oenocyte (Stringent) loom and h5ad file, encompassing a total of 506,660 cells. The Seurat (v4.2.2) package (https://doi.org/10.1016/j.cell.2021.04.048) was utilized for data analysis. Violin plots were generated using the “Vlnplot” function, the cell types are split by FCA.

### Statistical Tests

Statistical analysis of mating duration assay was described previously^36,37,48^. More than 50 males (naïve, experienced and single) were used for mating duration assay. Our experience suggests that the relative mating duration differences between naïve and experienced condition and singly reared are always consistent; however, both absolute values and the magnitude of the difference in each strain can vary. So, we always include internal controls for each treatment as suggested by previous studies^90^. Therefore, statistical comparisons were made between groups that were naïvely reared, sexually experienced and singly reared within each experiment. As mating duration of males showed normal distribution (Kolmogorov-Smirnov tests, p > 0.05), we used two-sided Student’s t tests. The mean ± standard error (s.e.m) (*\*\*\*\* = p < 0.0001, *** = p < 0.001, ** = p < 0.01, * = p < 0.05*). All analysis was done in GraphPad (Prism). Individual tests and significance are detailed in figure legends. Besides traditional *t*-test for statistical analysis, we added estimation statistics for all MD assays and two group comparing graphs. In short, ‘estimation statistics’ is a simple framework that—while avoiding the pitfalls of significance testing—uses familiar statistical concepts: means, mean differences, and error bars. More importantly, it focuses on the effect size of one’s experiment/intervention, as opposed to significance testing^91^. In comparison to typical NHST plots, estimation graphics have the following five significant advantages such as (1) avoid false dichotomy, (2) display all observed values (3) visualize estimate precision (4) show mean difference distribution. And most importantly (5) by focusing attention on an effect size, the difference diagram encourages quantitative reasoning about the system under study^92^. Thus, we conducted a reanalysis of all of our two group data sets using both standard *t* tests and estimate statistics. In 2019, the Society for Neuroscience journal eNeuro instituted a policy recommending the use of estimation graphics as the preferred method for data presentation^93^.

### Genotypes of flies used for experiments in this study

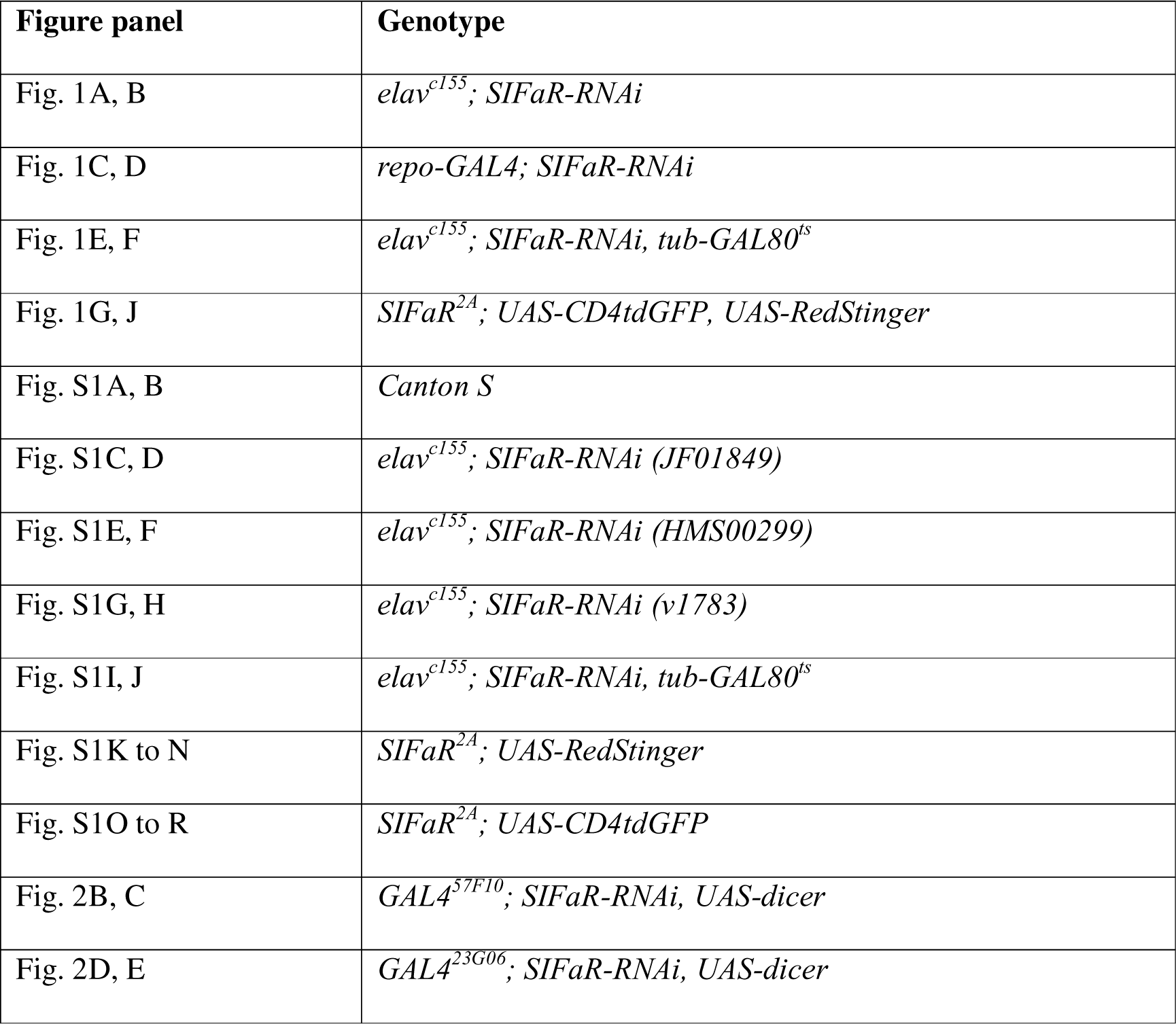
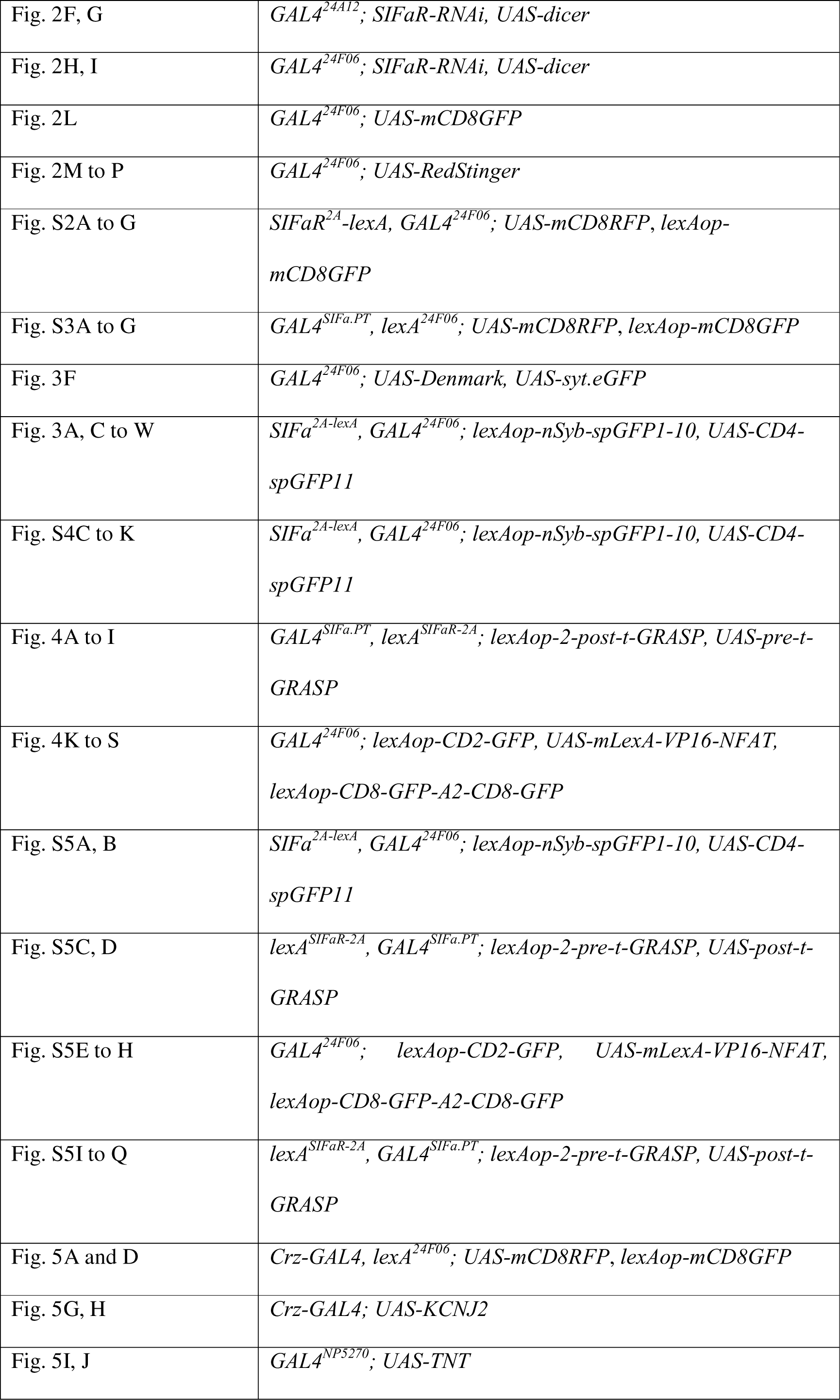
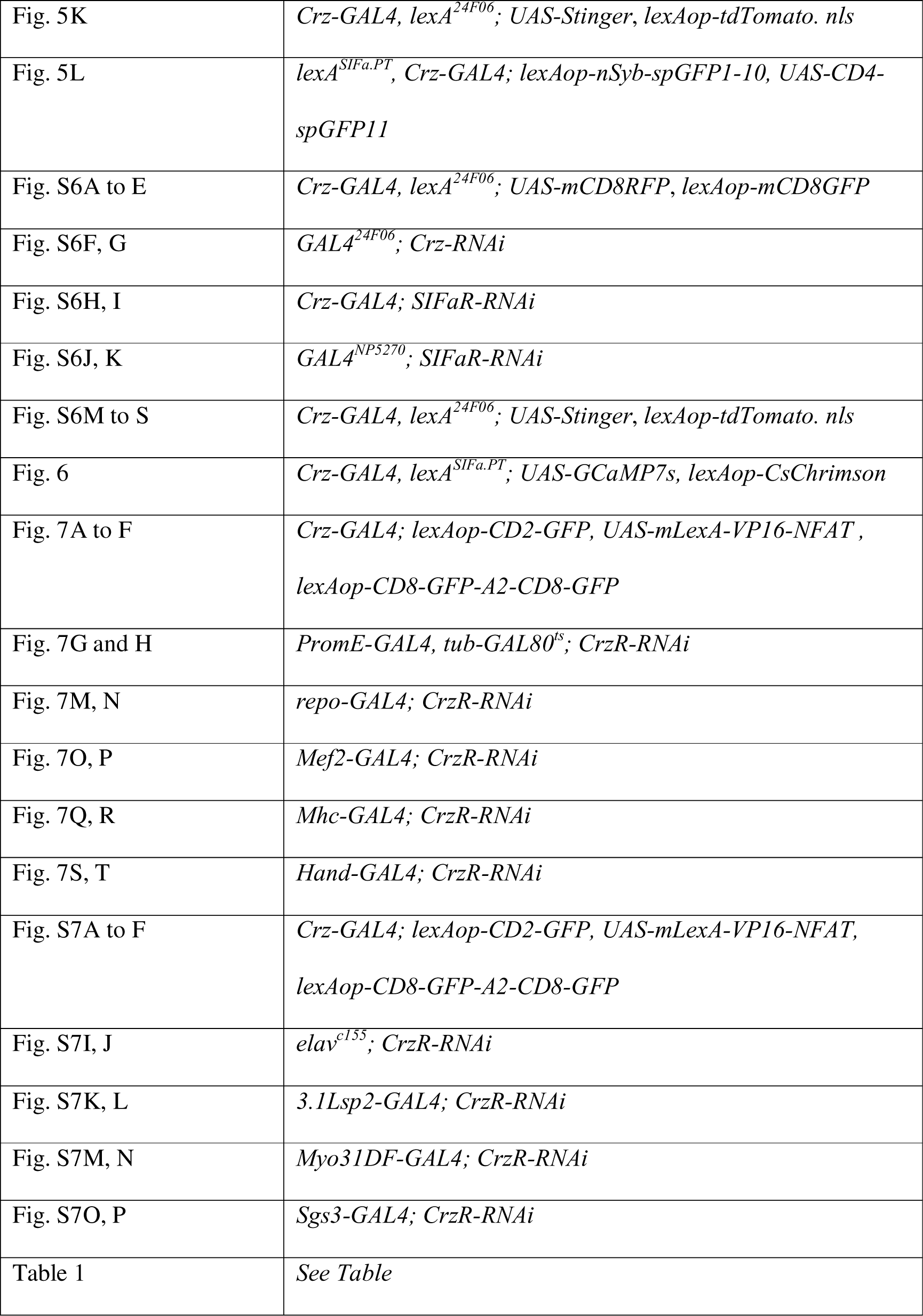

**Genotypes of flies used for experiments in this study.**
| Figure panel | Genotype |
| --- | --- |
| Fig. 1A, B | *elav<sup>c155</sup>; SIFaR-RNAi* |
| Fig. 1C, D | *repo-GAL4; SIFaR-RNAi* |
| Fig. 1E, F | *elav<sup>c155</sup>; SIFaR-RNAi, tub-GAL80<sup>ts</sup>* |
| Fig. 1G, J | *SIFaR<sup>2A</sup>; UAS-CD4tdGFP, UAS-RedStinger* |
| Fig. S1A, B | *Canton S* |
| Fig. S1C, D | *elav<sup>c155</sup>; SIFaR-RNAi (JF01849)* |
| Fig. S1E, F | *elav<sup>c155</sup>; SIFaR-RNAi (HMS00299)* |
| Fig. S1G, H | *elav<sup>c155</sup>; SIFaR-RNAi (v1783)* |
| Fig. S1I, J | *elav<sup>c155</sup>; SIFaR-RNAi, tub-GAL80<sup>ts</sup>* |
| Fig. S1K to N | *SIFaR<sup>2A</sup>; UAS-RedStinger* |
| Fig. S1O to R | *SIFaR<sup>2A</sup>; UAS-CD4tdGFP* |
| Fig. 2B, C | *GAL4<sup>57F10</sup>; SIFaR-RNAi, UAS-dicer* |
| Fig. 2D, E | *GAL4<sup>23G06</sup>; SIFaR-RNAi, UAS-dicer* |
| Fig. 2F, G | *GAL4<sup>24A12</sup>; SIFaR-RNAi, UAS-dicer* |
| Fig. 2H, I | *GAL4<sup>24F06</sup>; SIFaR-RNAi, UAS-dicer* |
| Fig. 2L | *GAL4<sup>24F06</sup>; UAS-mCD8GFP* |
| Fig. 2M to P | *GAL4<sup>24F06</sup>; UAS-RedStinger* |
| Fig. S2A to G | *SIFaR<sup>2A</sup>-lexA, GAL4<sup>24F06</sup>; UAS-mCD8RFP, lexAop-mCD8GFP* |
| Fig. S3A to G | *GAL4<sup>SIFa.PT</sup>, lexA<sup>24F06</sup>; UAS-mCD8RFP, lexAop-mCD8GFP* |
| Fig. 3F | *GAL4<sup>24F06</sup>; UAS-Denmark, UAS-syt.eGFP* |
| Fig. 3A, C to W | *SIFa<sup>2A-lexA</sup>, GAL4<sup>24F06</sup>; lexAop-nSyb-spGFP1-10, UAS-CD4-spGFP11* |
| Fig. S4C to K | *SIFa<sup>2A-lexA</sup>, GAL4<sup>24F06</sup>; lexAop-nSyb-spGFP1-10, UAS-CD4-spGFP11* |
| Fig. 4A to I | *GAL4<sup>SIFa.PT</sup>, lexA<sup>SIFaR-2A</sup>; lexAop-2-post-t-GRASP, UAS-pre-t-GRASP* |
| Fig. 4K to S | *GAL4<sup>24F06</sup>; lexAop-CD2-GFP, UAS-mLexA-VP16-NFAT, lexAop-CD8-GFP-A2-CD8-GFP* |
| Fig. S5A, B | *SIFa<sup>2A-lexA</sup>, GAL4<sup>24F06</sup>; lexAop-nSyb-spGFP1-10, UAS-CD4-spGFP11* |
| Fig. S5C, D | *lexA<sup>SIFaR-2A</sup>, GAL4<sup>SIFa.PT</sup>; lexAop-2-pre-t-GRASP, UAS-post-t-GRASP* |
| Fig. S5E to H | *GAL4<sup>24F06</sup>; lexAop-CD2-GFP, UAS-mLexA-VP16-NFAT, lexAop-CD8-GFP-A2-CD8-GFP* |
| Fig. S5I to Q | *lexA<sup>SIFaR-2A</sup>, GAL4<sup>SIFa.PT</sup>; lexAop-2-pre-t-GRASP, UAS-post-t-GRASP* |
| Fig. 5A and D | *Crz-GAL4, lexA<sup>24F06</sup>; UAS-mCD8RFP, lexAop-mCD8GFP* |
| Fig. 5G, H | *Crz-GAL4; UAS-KCNJ2* |
| Fig. 5I, J | *GAL4<sup>NP5270</sup>; UAS-TNT* |
| Fig. 5K | *Crz-GAL4, lexA<sup>24F06</sup>; UAS-Stinger, lexAop-tdTomato. nls* |
| Fig. 5L | *lexA<sup>SIFa.PT</sup>, Crz-GAL4; lexAop-nSyb-spGFP1-10, UAS-CD4-spGFP11* |
| Fig. S6A to E | *Crz-GAL4, lexA<sup>24F06</sup>; UAS-mCD8RFP, lexAop-mCD8GFP* |
| Fig. S6F, G | *GAL4<sup>24F06</sup>; Crz-RNAi* |
| Fig. S6H, I | *Crz-GAL4; SIFaR-RNAi* |
| Fig. S6J, K | *GAL4<sup>NP5270</sup>; SIFaR-RNAi* |
| Fig. S6M to S | *Crz-GAL4, lexA<sup>24F06</sup>; UAS-Stinger, lexAop-tdTomato. nls* |
| Fig. 6 | *Crz-GAL4, lexA<sup>SIFa.PT</sup>; UAS-GCaMP7s, lexAop-CsChrimson* |
| Fig. 7A to F | *Crz-GAL4; lexAop-CD2-GFP, UAS-mLexA-VP16-NFAT, lexAop-CD8-GFP-A2-CD8-GFP* |
| Fig. 7G and H | *PromE-GAL4, tub-GAL80<sup>ts</sup>; CrzR-RNAi* |
| Fig. 7M, N | *repo-GAL4; CrzR-RNAi* |
| Fig. 7O, P | *Mef2-GAL4; CrzR-RNAi* |
| Fig. 7Q, R | *Mhc-GAL4; CrzR-RNAi* |
| Fig. 7S, T | *Hand-GAL4; CrzR-RNAi* |
| Fig. S7A to F | *Crz-GAL4; lexAop-CD2-GFP, UAS-mLexA-VP16-NFAT, lexAop-CD8-GFP-A2-CD8-GFP* |
| Fig. S7I, J | *elav<sup>c155</sup>; CrzR-RNAi* |
| Fig. S7K, L | *3.1Lsp2-GAL4; CrzR-RNAi* |
| Fig. S7M, N | *Myo31DF-GAL4; CrzR-RNAi* |
| Fig. S7O, P | *Sgs3-GAL4; CrzR-RNAi* |
| Table 1 | *See Table* |

## ACKNOWLEDGMENTS

We thank Dr. Jan A. Veenstra (University of Bordeaux) for sharing SIFa^PT^-GAL4 driver, Dr. Young-Joon Kim (GIST) for kindly sharing UAS-SIFaR fly strain, Drs. Yuh Nung Jan and Lily Yeh Jan (UCSF, USA) for helpful comments and support on this paper. We are also very appreciative to the colleagues who supplied us with several fly strains: Dr. Wei Zhang (Tsinghua University), Fang Guo (Zhejiang University), and Dr. Yufeng Pan (Southeast University) and Drs. Young-Joon Kim and Sung-Eun Yoon (Korea Drosophila Resource Center, KDRC). This research was supported a University of Ottawa Startup grant 602496 to WJK, Startup funds from HIT Center for Life Science to WJK, a University of Ottawa Interdisciplinary Research Group Funding Opportunity (IRGFO stream 1 and 2) grants 148101 and 148747 to WJK, a Natural Sciences and Engineering Research Council of Canada (NSERC) Discovery grant (reference: 211406) to WJK, a University of Ottawa Brain and Mind Research Institute/Center for Neural Dynamics Open call project grant 150950 to WJK, a Mitacs Globalink Research Internship Program grant 17268 to WJK. This research was also supported by the Brain Pool Program of the National Research Foundation in Korea grant ZYM5041911 to WJK, Burroughs Wellcome Fund Collaborative Research Travel Grants (reference: 1017486) to WJK and a NVIDIA Academic Hardware Grant Program to WJK. The funders had no role in study design, data collection and analysis, decision to publish, or preparation of the manuscript. SGL received salary from the ‘University of Ottawa Startup grant to WJK’ and HM from the ‘Startup funds from HIT Center for Life Science to WJK’.

## AUTHOR CONTRIBUTIONS

**Conceptualization:** Woo Jae Kim.

**Data curation:** Tianmu Zhang, Yutong Song, Zekun Wu, Kyle Wong, Justine Schweizer, Woo Jae Kim.

**Formal analysis:** Tianmu Zhang, Zekun Wu, Yutong Song, Wenjing Li, Yanying Sun, Xiaoli Zhang, Kyle Wong, Justine Schweizer, Khoi-Nguyen Ha Nguyen, Alex Kwan, and Woo Jae Kim.

**Funding acquisition:** Woo Jae Kim.

**Investigation:** Woo Jae Kim.

**Methodology:** Woo Jae Kim.

**Project administration:** Woo Jae Kim.

**Resources:** Woo Jae Kim.

**Supervision:** Woo Jae Kim.

**Validation:** Tianmu Zhang, Zekun Wu, Woo Jae Kim.

**Visualization:** Zekun Wu, Woo Jae Kim.

**Writing – original draft:** Woo Jae Kim.

**Writing – review & editing:** Tianmu Zhang, Woo Jae Kim.

## DECLARATION OF GENERATIVE AI AND AI-ASSISTED TECHNOLOGIES IN THE WRITING PROCESS

During the creation of this work, the author(s) utilized QuillBot to rephrase English sentences, verify English grammar, and detect plagiarism, as none of the authors of this paper are native English speakers. After using this tool/service, the author(s) reviewed and edited the content as needed and take(s) full responsibility for the content of the publication.

## SUPPLEMENTAL FIGURE TITLES AND LEGENDS

**Figure S1.**
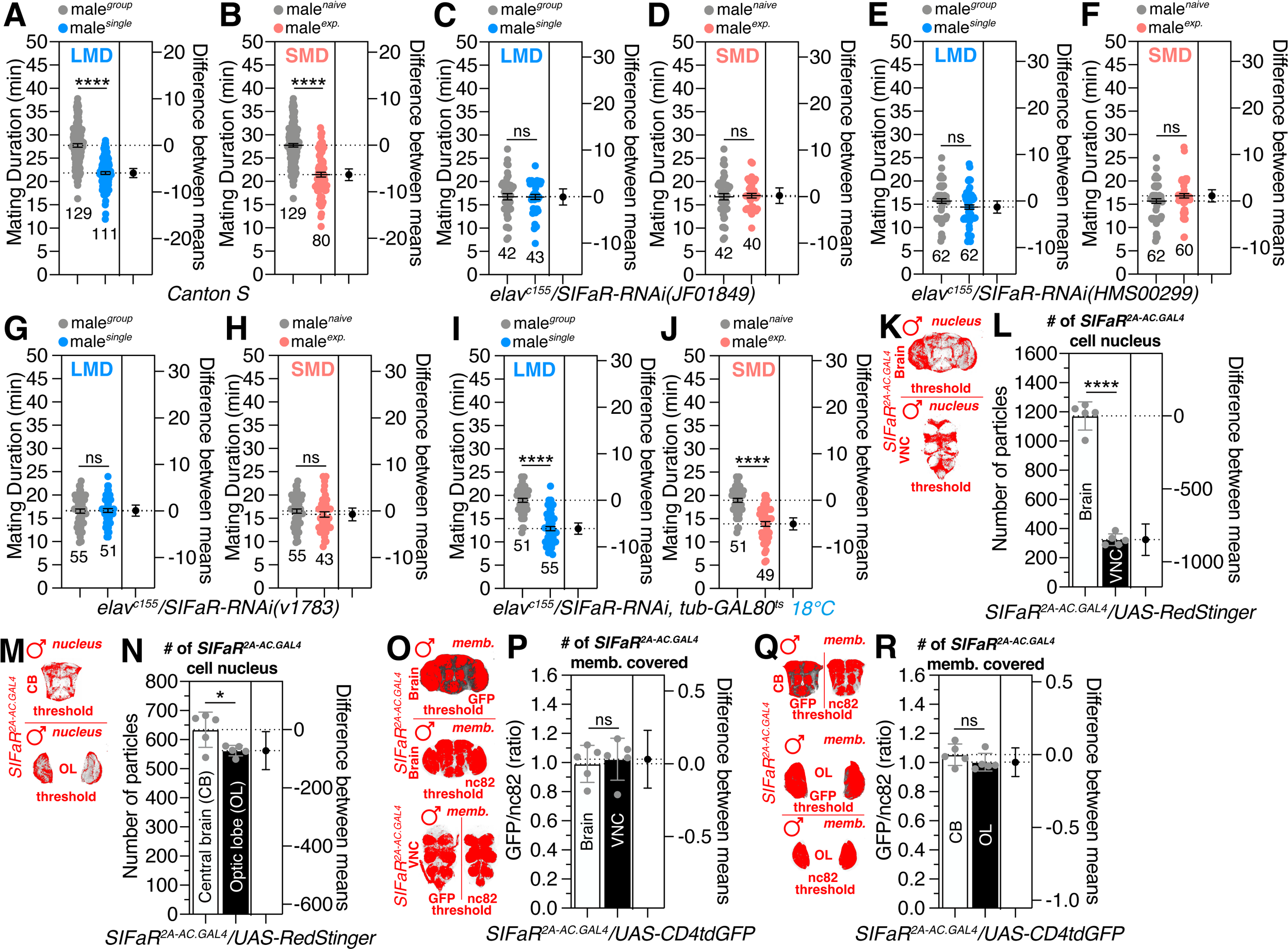
Screening of *SIFaR-RNAi* lines and quantification of *SIFaR^2A^* expression in CNS. Related to Figure 1. (A-B) LMD and SMD assays of *Conton-S* (WT). (C-J) LMD and SMD assays for *elav^c155^*–mediated knockdown of SIFaR *via* (C-D) *SIFaR-RNAi (JF01849),* (E-F) *SIFaR-RNAi (HMS00299)* and (G-H) *SIFaR-RNAi (HMS00299)*. (I-J) LMD and SMD assays for *elav^c155^*–mediated knockdown of SIFaR *via SIFaR-RNAi* together with *tub-GAL80^ts^* in 18. (K-L) Quantification of cell number of male fly brain and VNC expressing *SIFaR^2A^* driver together with *UAS-Redstinger*. (M-N) Quantification of cell number in male fly CB and OL expressing *SIFaR^2A^* driver together with *UAS-Redstinger*. (O-R) Quantification of GFP fluorescence in male fly brain expressing *SIFaR^2A^* driver together with *UAS-CD4tdGFP*.

**Figure S2.**
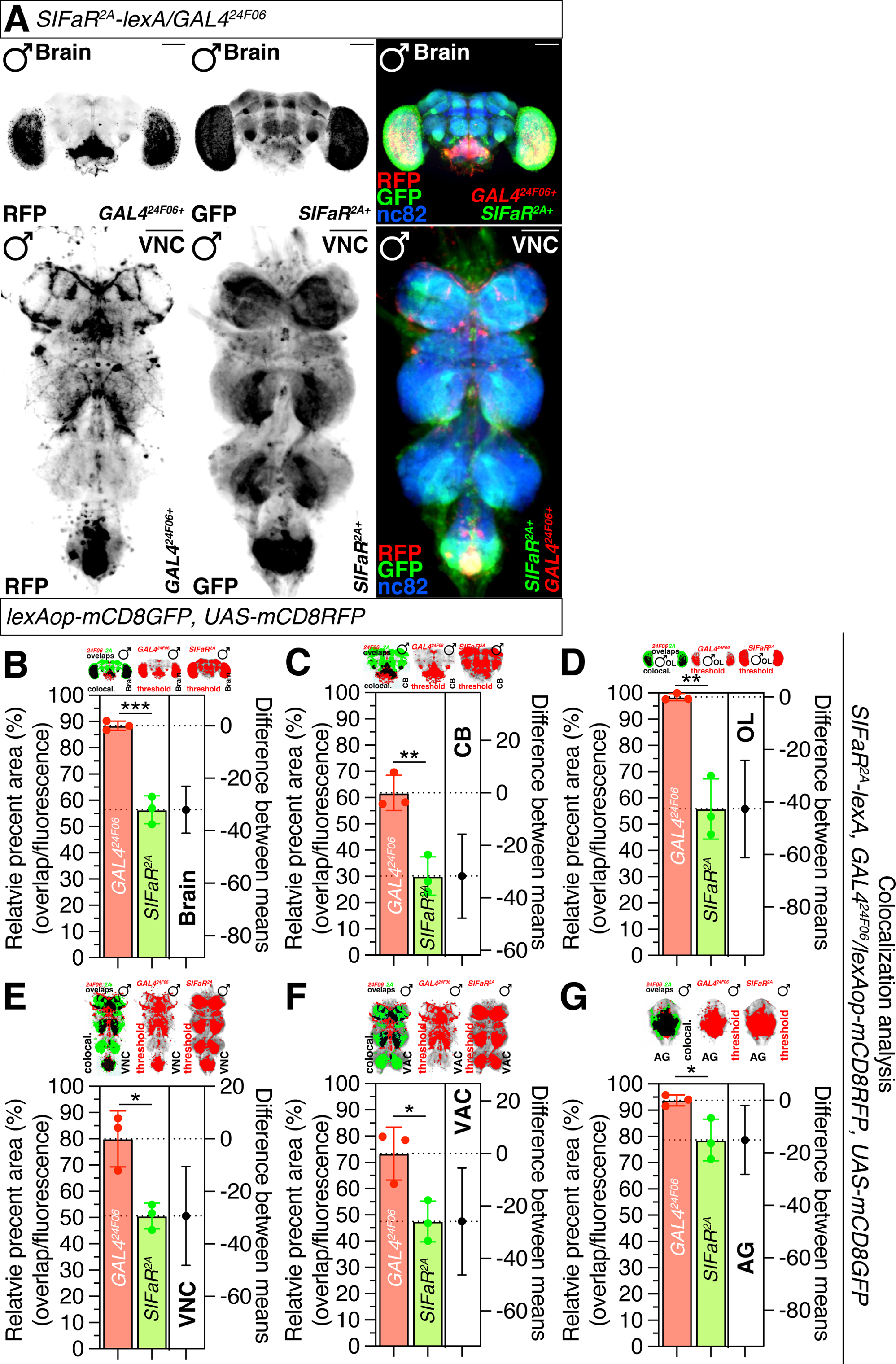
Colocalization analysis between *GAL4^24F06^* and *SIFaR^2A^-lexA*. Related to Figure 2. (A) Male flies brain expressing *SIFaR^2A^-lexA* and *GAL4^24F06^* drivers together with *UAS-mCD8RFP* and *lexAop-mCD8GFP* were immunostained with anti-GFP (green), anti-DsRed (red) and anti-nc82 (blue) antibodies. Scale bars represent 100 μm. Boxes indicate the magnified regions of interest presented in the bottom panels. The left two panels are presented as a grey scale to clearly show the axon projection patterns of neurons in brain and VNC labeled by *SIFaR^2A^ -lexA* and *GAL4^24F06^* driver. (B-G) Colocalization analysis of GFP and RFP staining, normalized to total GFP and RFP areas. See the MATERIALS AND METHODS for a detailed description of the colocalization analysis used in this study.

**Figure S3.**
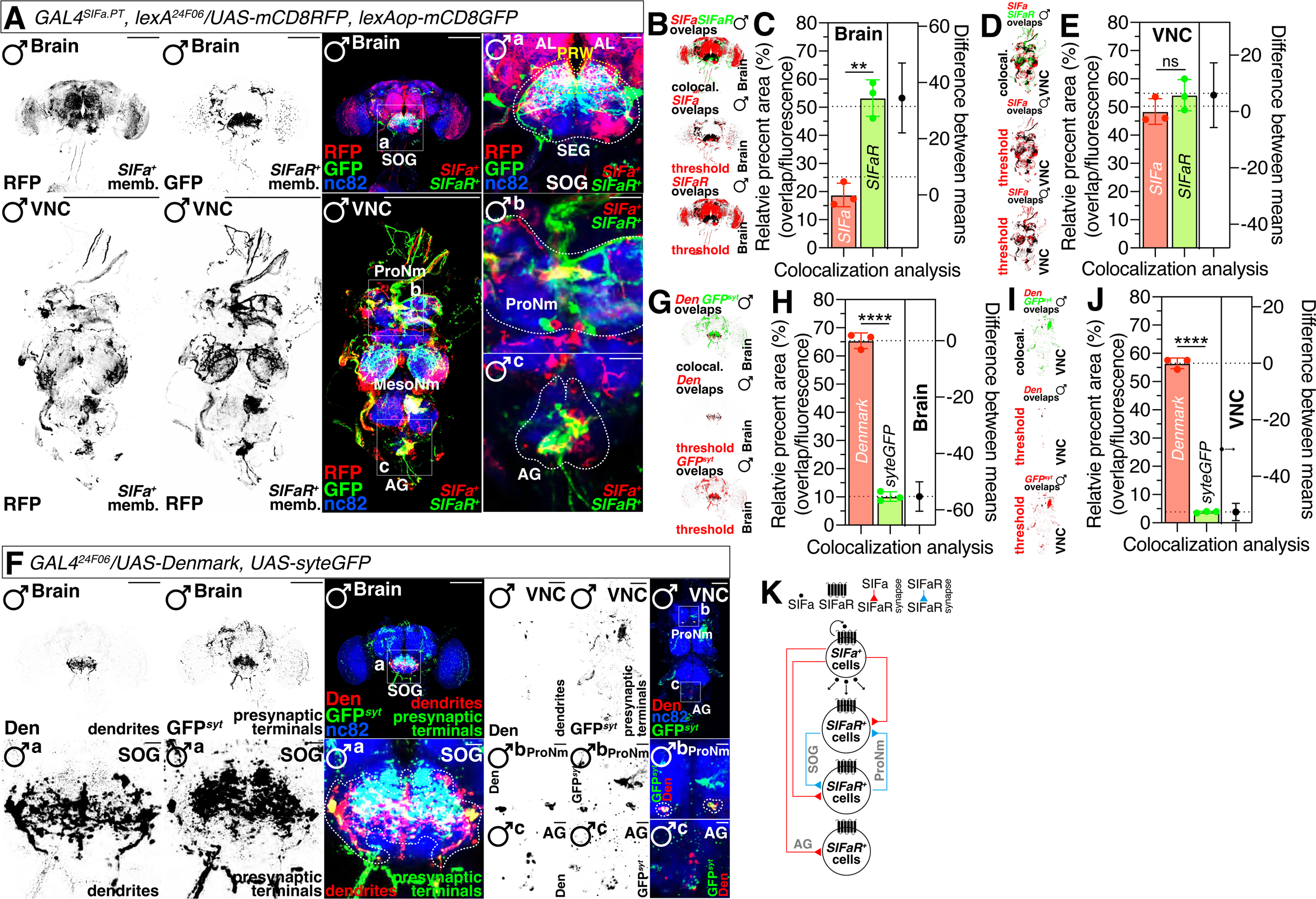
*GAL4^SIFa.PT^* and *lexA^24F06^* form strong synapses and *SIFaR^24F06^*-positive neurons interconnected in specific regions in CNS. Related to Figure 2. (A) Male flies brain expressing *GAL4^SIFa.PT^* and *lexA^24F06^* drivers together with *UAS-mCD8RFP* and *lexAop-mCD8GFP* were immunostained with anti-GFP (green), anti-DsRed (red) and anti-nc82 (blue) antibodies. Boxes and dashed circles indicate the magnified regions of interest presented in the bottom panels. The left two panels are presented as a grey scale to clearly show the axon projection patterns of neurons labeled by *GAL4^SIFa.PT^* and *lexA^24F06^* driver. Scale bars represent 100 μm in brain and VNC panels, and 25 μm in other panels. (B-E) Colocalization analysis of GFP and RFP staining, normalized to total GFP and RFP areas. See the MATERIALS AND METHODS for a detailed description of the colocalization analysis used in this study. (F) Male flies brain expressing *GAL4^24F06^* drivers together with *UAS-Denmark* and *UAS-syt.eGFP* were immunostained with anti-GFP (green), anti-DsRed (red) and anti-nc82 (blue) antibodies. Boxes indicate the magnified regions of interest presented in the bottom panels. Dashed circles indicate the neurons where *GAL4^24F06^* form interconnected networks. The left two panels are presented as a grey scale to clearly show the dendritic and synaptic terminal of neurons labeled by *GAL4^24F06^* driver. Scale bars represent 100 μm in brain panels, 50 μm in VNC panels, 10 μm in SOG panels, and 25 μm in ProNm and AG panels. (G-J) Colocalization analysis of GFP and RFP staining, normalized to total GFP and RFP areas. See the MATERIALS AND METHODS for a detailed description of the colocalization analysis used in this study. (K) Diagram of interconnection between *GAL4^SIFa.PT^* and *lexA^24F06^* neurons along with interconnected networks formed by *GAL4^24F06^*.

**Figure S4.**
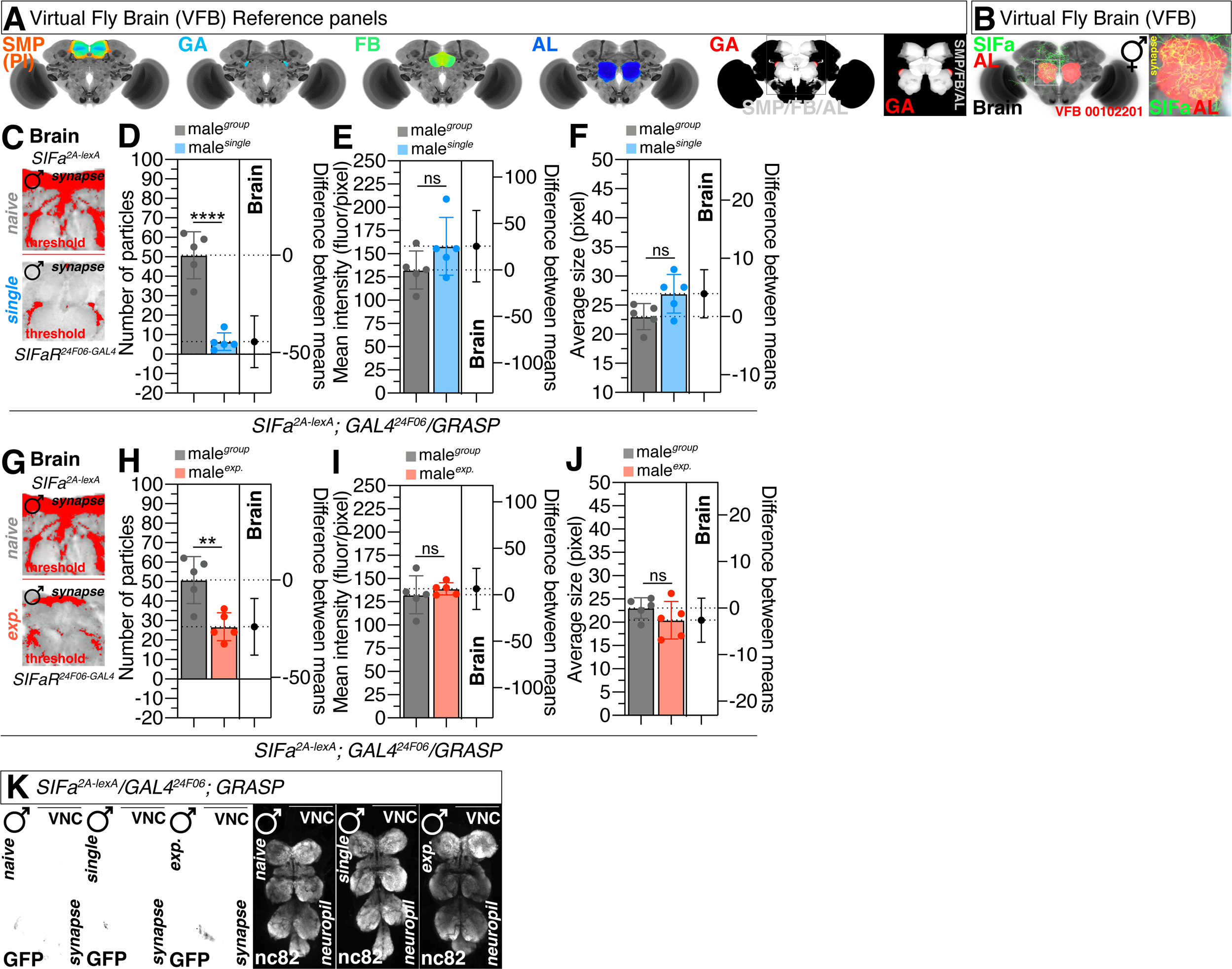
*SIFa^2A-lexA^* and *GAL4^24F06^* form synapses only in brain region. Related to Figure 3. (A) Virtual Fly Brain (VFB) visualization of SMP, GA, FB and AL brain regions in *Drosophila*. In the last two panels, the grey area represents the overlapping region of SMP/FB/AL, and the red area represents GA. (B) Co-locolization between SIFa and AL in Virtual Fly Brain (VFB). White boxes indicate the magnified regions of interest presented in the right panel. The yellow area represents synaptic connections formed by *SIFa* (green) and AL (red). (C) The synaptic interactions visualized utilizing the GRASP system in naïve and single male flies. The GFP fluorescence was processed using ImageJ software, where a threshold function was applied to distinguish fluorescence from the background. Synaptic transmission occurs from *SIFa^2A-lexA^* to *GAL4^24F06^*. (D) Quantification of synaptic puncta formed between *SIFa^2A-lexA^* and *GAL4^24F06^* in brain between naïve and single male flies. The synaptic interactions were visualized utilizing the GRASP system in male flies. (E) Quantification of synapses intensity formed between *SIFa^2A-lexA^* and *GAL4^24F06^* in brain between naïve and single male flies. The synaptic interactions were visualized utilizing the GRASP system in male flies. (E) Quantification of average synapse size formed between *SIFa^2A-lexA^* and *GAL4^24F06^* in brain between naïve and single male flies. The synaptic interactions were visualized utilizing the GRASP system in male flies. (G) The synaptic interactions visualized utilizing the GRASP system in naïve and experienced male flies. The GFP fluorescence was processed using ImageJ software, where a threshold function was applied to distinguish fluorescence from the background. Synaptic transmission occurs from *SIFa^2A-lexA^* to *GAL4^24F06^* (H) Quantification of synaptic puncta formed between *SIFa^2A-lexA^* and *GAL4^24F06^* in brain between naïve and single male flies. The synaptic interactions were visualized utilizing the GRASP system in male flies. (I) Quantification of synapses intensity formed between *SIFa^2A-lexA^* and *GAL4^24F06^* in brain between naïve and experienced male flies. The synaptic interactions were visualized utilizing the GRASP system in male flies. (J) Quantification of average synapse size formed between *SIFa^2A-lexA^* and *GAL4^24F06^* in brain between naïve and experienced male flies. The synaptic interactions were visualized utilizing the GRASP system in male flies. (K) No synapses were formed between *SIFa^2A-lexA^* and *GAL4^24F06^* in VNC. Scale bars represent 100 μm

**Figure S5.**
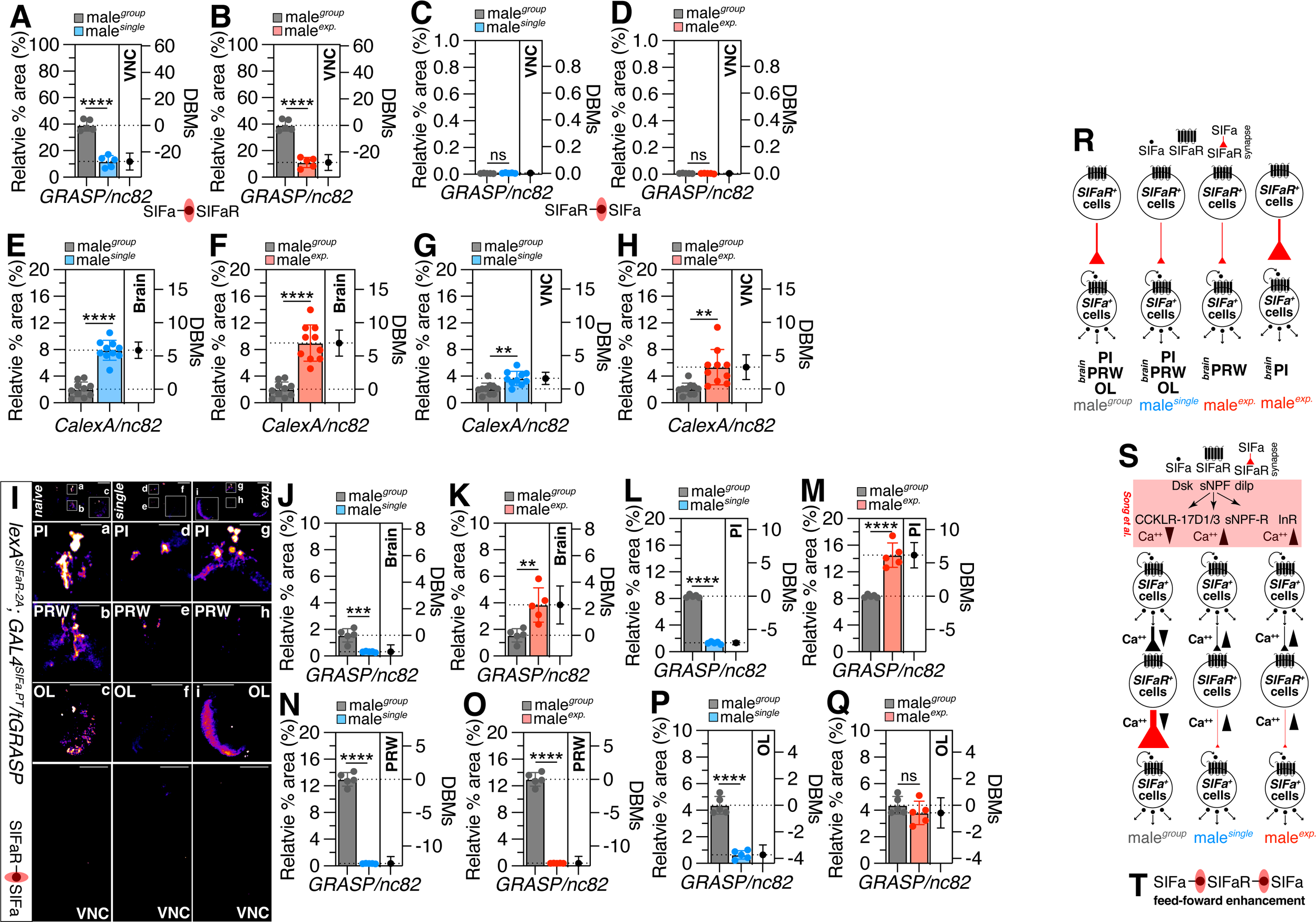
Social context modulates both the formation of synapses between *GAL4^SIFa.PT^* and *lexA^24F06^* neurons in VNC and the calcium-dependent activity of *SIFaR*. Related to Figure 4. (A-B) Quantification of synaptic relative area formed between *GAL4^SIFa.PT^* and *lexA^24F06^* in VNC between (A) naïve and single male flies; (B) naïve and experienced male flies. The synaptic interactions were visualized utilizing the tGRASP system in naïve, single and experienced male flies. Synaptic transmission occurs from *GAL4^SIFa.PT^* to *lexA^24F06^*. (C-D) Quantification of synaptic relative area formed between *lexA^SIFaR-2A^* and *GAL4^SIFa.PT^* in VNC between (C) naïve and single male flies; (D) naïve and experienced male flies. The synaptic interactions were visualized utilizing the tGRASP system in naïve, single and experienced male flies. Synaptic transmission occurs from *lexA^SIFaR-2A^* to *GAL4^SIFa.PT^*. (E-H) Quantification of GFP fluorescence in brain and VNC between (E, G) naïve and single male flies. The same quantification was performed for the GFP fluorescence in these regions between naïve and experienced male flies. The GFP fluorescence was normalized relative to the fluorescence of the nc82. The conditions of flies are described above: naïve, naïve male flies; single, singly reared male flies; exp., male flies with sexual experience. (I) tGRASP assay for *GAL4^SIFa.PT^* and *lexA^SIFaR-2A^* in whole brain, PI, PRW, OL and VNC region of male flies. Male flies expressing *GAL4^SIFa.PT^* and *lexA^SIFaR-2A^* and *LexAop-2-pre-t-GRASP, UAS-post-t-GRASP* were dissected after 5 days of growth. Brains of male flies were immunostained with anti-GFP (green) and anti-nc82 (blue) antibodies. GFP is pseudo-colored as “red hot”. Boxes indicate the magnified regions of interest presented in the bottom panels. Scale bars represent 100 μm in brain and VNC panels, 25μm in PI and PRW panels, and 50 μm in OL panels. Synaptic transmission occurs from *lexA^SIFaR-2A^* to *GAL4^SIFa.PT^*. (J-Q) Quantification of synaptic relative area formed between *GAL4^SIFa.PT^* and *lexA^SIFaR-2A^* in (J) brain, (L) PI, (N) PRW and (P) OL between naïve and single male flies. The same quantification was performed for the relative synaptic area in these regions between naïve and experienced male flies. The synaptic interactions were visualized utilizing the tGRASP system in naïve, single and experienced male flies. (R) Diagram of differential SIFa-SIFaR signaling across various regions of the CNS in male *Drosophila melanogaster*, contingent upon diverse social contexts. (S) Schematic representation of neuronal circuits modulating CNS internal states through feed-forward augmentation in response to social contexts. (T) Feed-forward enhancement circuits of *SIFa-SIFaR-SIFa*.

**Figure S6.**
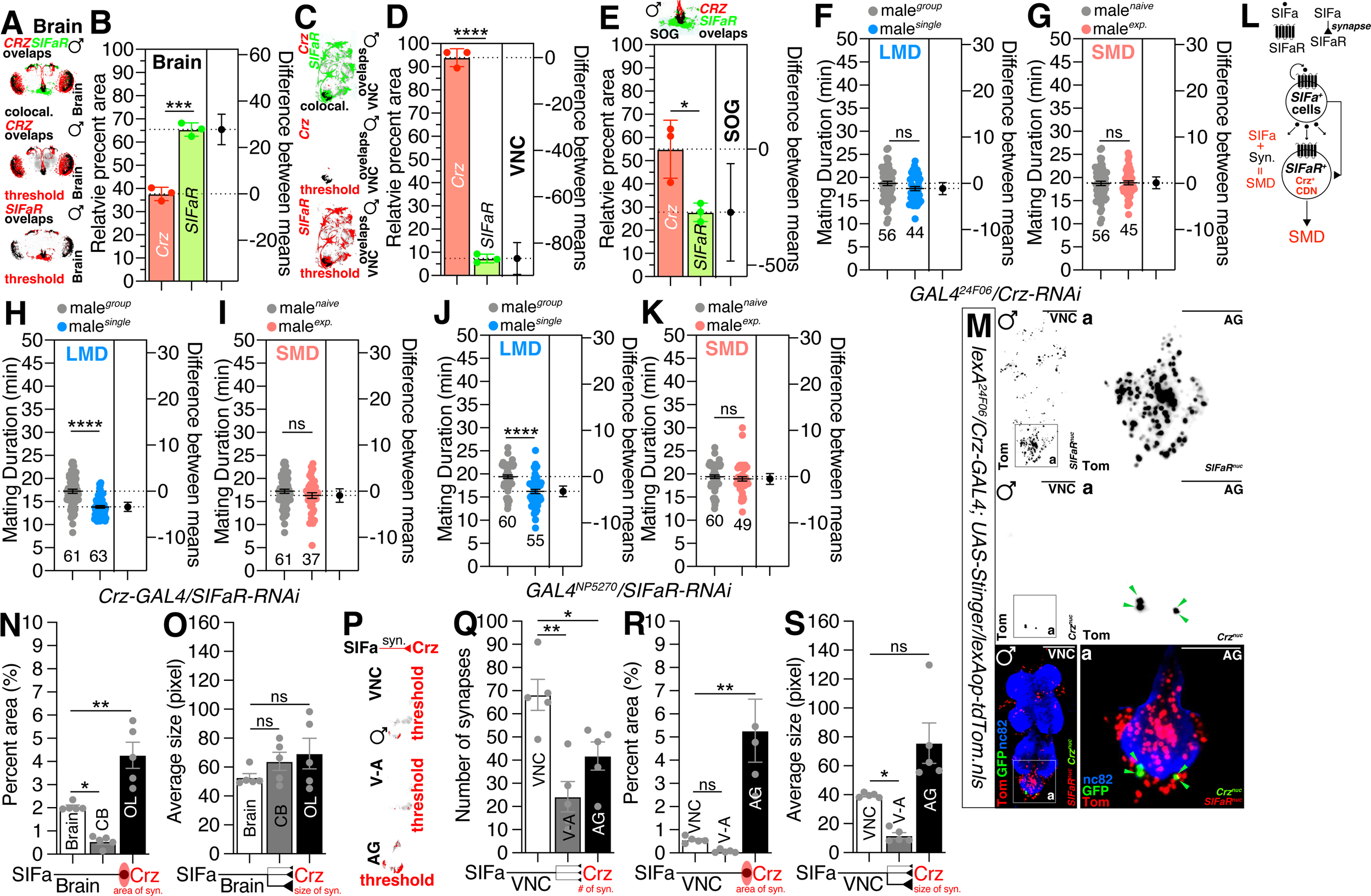
CDN and *Crz* signaling *via SIFaR* predominantly shapes SMD behavior. Related to Figure 5. (A-E) Colocalization analysis of GFP and RFP staining, normalized to total GFP and RFP areas. See the MATERIALS AND METHODS for a detailed description of the colocalization analysis used in this study. (F-G) LMD and SMD assays for *GAL4^24F06^* mediated knockdown of Crz *via Crz-RNAi*. (H-I) LMD and SMD assays for *Crz-GAL4* mediated knockdown of SIFaR *via SIFaR-RNAi*. (J-K) LMD and SMD assays for *GAL4^NP5270^* mediated knockdown of SIFaR *via SIFaR-RNAi*. (L) Diagram of the CDN and Crz signaling pathway through SIFa-SIFaR signaling in male *Drosophila melanogaster* in SMD behavior. (M) Male flies VNC expressing *Crz-GAL4* and *lexA^24F06^* drivers together with *UAS-Stinger* and *lexAop-tdTomato.nls* were immunostained with anti-GFP (green), anti-DsRed (red) and anti-nc82 (blue) antibodies. Scale bars represent 100 μm in VNC panels and 50 μm in AG panels. Boxes indicate the magnified regions of interest presented in the bottom panels. The upper panels are presented as a grey scale to clearly show the nucleus in the adult VNC and AG labeled by *Crz-GAL4* and *lexA^24F06^* driver. Green arrows indicate *Crz^+^* nucleus. (N) Quantification of synaptic relative area formed between *SIFa^PT^-lexA* and *Crz-GAL4* in brain, CB and OL in male flies. The synaptic interactions were visualized utilizing the tGRASP system in male flies. Synaptic transmission occurs from *SIFa^PT^-lexA* to *Crz-GAL4*. (O) Quantification of average synapse size formed between *SIFa^PT^-lexA* and *Crz-GAL4* in brain, CB and OL in male flies. The synaptic interactions were visualized utilizing the GRASP system in male flies. Synaptic transmission occurs from *SIFa^PT^-lexA* to *Crz-GAL4*. (P) The GFP fluorescence was processed using ImageJ software, where a threshold function was applied to distinguish fluorescence from the background. (Q) Quantification of synaptic puncta formed between *SIFa^PT^-lexA* and *Crz-GAL4* in VNC, V-A and AG in male flies. The synaptic interactions were visualized utilizing the tGRASP system in male flies. Synaptic transmission occurs from *SIFa^PT^-lexA* to *Crz-GAL4*. (R) Quantification of synaptic relative area formed between *SIFa^PT^-lexA* and *Crz-GAL4* in VNC, V-A and AG in male flies. The synaptic interactions were visualized utilizing the tGRASP system in male flies. Synaptic transmission occurs from *SIFa^PT^-lexA* to *Crz-GAL4*. (S) Quantification of average synapse size formed between *SIFa^PT^-lexA* and *Crz-GAL4* in VNC, V-A and AG in male flies. The synaptic interactions were visualized utilizing the GRASP system in male flies. Synaptic transmission occurs from *SIFa^PT^-lexA* to *Crz-GAL4*.

**Figure S7.**
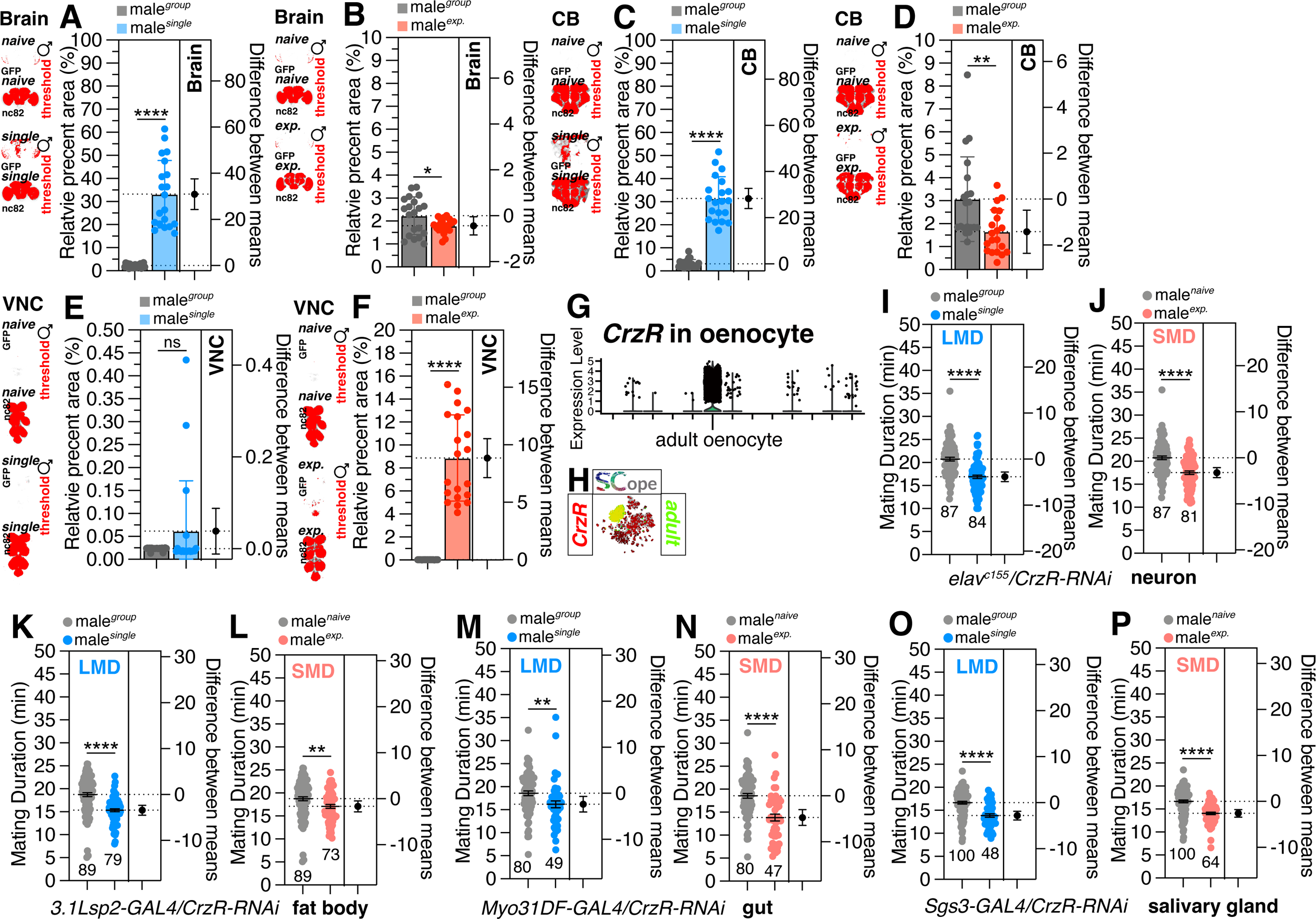
Non-neuronal cells required for SIFa-SIFaR/Crz-CrzR signaling in interval timing behaviors. Related to Figure 6. (A-F) Quantification of GFP fluorescence in (A) brain, (C) CB and (E) VNC between naïve and single male flies. The same quantification was performed for the GFP fluorescence in these regions between naïve and experienced male flies. The GFP fluorescence was normalized relative to the fluorescence of the nc82. The conditions of flies are described above: naïve, naïve male flies; single, singly reared male flies; exp., male flies with sexual experience. The small panels are presented as a red scale to show the GFP fluorescence marked by threshold function of ImageJ. (G) Dot plot depicting the expression levels of *CrzR* in ‘adult oenocyte’ as annotated by the Fly Cell Atlas (FCA) within the oenocyte tissue. The gene expression levels for individual cells were normalized using the ‘LogNormalize’ method with a scale factor of 10,000, followed by scaling of all genes. (H) Single-cell RNA sequencing (SCOPE scRNA-seq) datasets reveal cell clusters colored by expression of *CrzR* (red) and adult oenocyte (green) in oenocyte. (I-P) LMD and SMD assays for *elav^c155^* (neuron)*, 3.1Lsp2-GAL4* (fat body)*, Myo31DF-GAL4* (gut) and *Sgs3-GAL4* (salivary gland)-mediated knockdown of CrzR *via CrzR-RNAi*.

